# Harnessing Microbial Tools: *Escherichia coli* as a Vehicle for Neuropeptide Functional Analysis in *Caenorhabditis elegans*

**DOI:** 10.1101/2025.03.03.641308

**Authors:** Elizabeth M. DiLoreto, Shruti Shastry, Emily J. Leptich, Douglas K. Reilly, Rachel N. Arey, Jagan Srinivasan

## Abstract

Animals respond to changes in their environment and internal states via neuromodulation. Neuropeptides modulate neural circuits with flexibility because one gene can produce either multiple copies of the same neuropeptide or different neuropeptides. However, with this architectural complexity, the function of discrete and active neuropeptides is muddled. Here, we design a genetic tool that facilitates functional analysis of individual peptides. We engineered *Escherichia coli* bacteria to express active peptides, fed loss-of-function *Caenorhabditis elegans*, and rescued the activity of genes with varying lengths and functions: *pdf-1, flp-3, ins-6*, and *ins-22*. Some peptides were functionally redundant, while others exhibit unique and previously uncharacterized functions. We postulate our rescue-by-feeding approach can elucidate the functional landscape of neuropeptides, identifying the circuits and complex peptidergic pathways that regulate different behavioral and physiological processes.

**Article summary:** Studying individual neuropeptides opens new avenues for exploring neuromodulation at a finer resolution. The researchers developed a method to create DNA vectors that encode an endogenous peptide sequence flanked by sequences containing dibasic endopeptidase cleavage sites in *Caenorhabditis elegans*. The researchers transformed these vectors into bacteria and fed them to *C. elegans*, which restored wildtype behavior in neuropeptide loss-of-function mutants. The researchers also discovered that neuropeptides from the same gene perform distinct functions, a research area more ready to explore using the presented technology.

**Graphical Abstract:** Created in BioRender. DiLoreto, E. (2025) https://BioRender.com/ypbrtlk

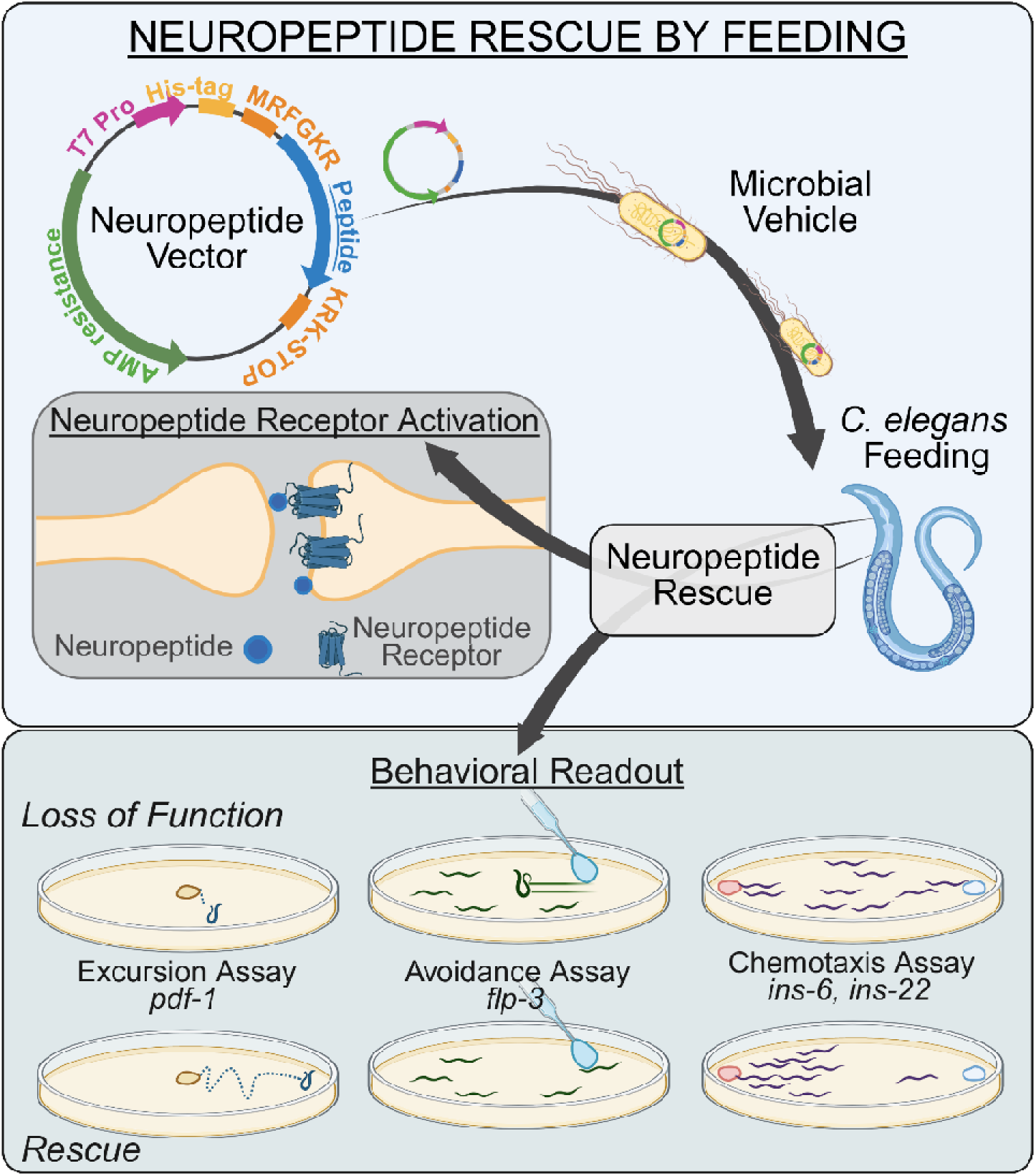

## Introduction

The modulation of neural circuits is essential for life, as appropriate responses to environmental stimuli depend on precise neural tuning. Neuropeptides, which are relatively short amino acid chains (less than 100 amino acids), play a crucial role in this process by regulating neural circuits while also functioning as neurotransmitters or neurohormones [1, 2]. They are widely used across the animal kingdom, even in organisms that possess only secretory cells without neurons [3]. Neuropeptides can also have conserved structure across species while serving a variety of functions. Oxytocin-like and vasopressin-like peptides are ancient neurohormones that regulate social attachment, lactation, and blood pressure in mammals [4–9]. However, in the Echinoderm sea star, *Asterias rubens*, the vasopressin-like and oxytocin-like neuropeptide ortholog, asterotocin, serves instead to relax muscles in the cardiac stomach during fictive feeding[9]. The *Caenorhabditis elegans* vasopressin ortholog, nematocin, interacts with serotonin and dopamine signaling to modulate gustatory associative memory and male mating behaviors [4, 10].

Neuropeptides act in conjunction with neurotransmitters, typically binding to G-protein coupled receptors, where they modulate synaptic activity more slowly and over a longer duration than neurotransmitters alone [2, 11]. Multiple modulators enable neurons to serve distinct functions within specific neural circuits [12]. Beyond the nervous system, neuropeptides also regulate immune and endocrine functions [13, 14]. In humans, there are varying reports of 90 or 131 neuropeptide precursor genes, yet all studies agree that there are likely more neuropeptides in the 900 other peptide genes [15–18]. Post-transcriptional modifications can give rise to an even greater number of functional neuropeptides, numbering in the thousands [15–19]. This complexity is even seen in a “simpler” model organism, the roundworm *Caenorhabditis elegans*. *C. elegans* is a microscopic nematode that displays robust behaviors driven by just over 300 neurons [20, 21]. The *C. elegans* genome encodes three classes of neuropeptides-FMRFamide-like peptides (FLP), insulin-like peptides (INS), and non-FLP/insulin neuropeptide-like peptides (NLP) [22–24]- and encodes over 300 individual neuropeptides through 160 identified genes that modulate the functional connectome [1, 19, 25, 26]. The complexity of the neuropeptide genome, combined with extra-synaptic neuropeptide signaling, makes elucidating the role of individual peptides difficult, as canonical studies often rely on null mutations and transgenic rescues [27–30]. In particular, full gene rescue restores complete preproproteins and makes discrimination of discrete peptide function difficult [1, 31]. Previous approaches to study individual endogenous neuropeptide function involved using synthetic peptides applied by soaking the nervous system of the worm [32] or injecting the peptide into the worm [33]. The use of synthetic peptides can be costly as pure peptide must be synthesized, often outsourced, for every neuropeptide experiment. Interestingly, feeding of exogenous peptides via *E. coli* has proven successful in manipulating *C. elegans* biological function [34]. However, this type of *E. coli* feeding of peptides has not been accomplished with endogenous neuropeptides.

Here, we report an approach that enables the manipulation of neuropeptide activity by feeding neuropeptide macromolecules produced by bacteria, allowing for the simultaneous delivery of individual neuropeptides to large populations of animals. Our gain-of-function approach of feeding bacteria containing the neuropeptide macromolecules (mRNA and peptides) provides ease of application and cost-saving measures to deliver endogenous peptides to *C. elegans.* Neuropeptide rescue by feeding, outlined here, expands on RNA interference (RNAi) feeding paradigms, which have been successfully used to test gene function [35, 36], wherein a plasmid encoding the RNA of interest is driven by isopropyl-β-D-thiogalactoside (IPTG)-induction. Whereas RNAi employs paired T7 promoters facing one another to produce double-stranded RNA, our protocol utilizes a single T7 promoter to generate mRNA encoding the peptide of interest. Once the neuropeptide vector is assembled and transformed into *E. coli*, delivery of the peptide simply requires feeding the worms. This method allows for temporal control of peptide delivery, animals can be fed at any life stage, and the number of animals treated is readily flexible.

The paradigm rescues individual, processed neuropeptides, leveraging the genetic amenability of the *C. elegans* food source, *Escherichia coli*, to circumvent the need for transgenic development and enable high-throughput rescue and elucidation of individual neuropeptide function [37]. We demonstrate the application of this paradigm by rescuing behaviors driven by neuropeptides synthesized from *pdf-1*, *ins-6*, and *ins-22*, and expand upon the previously established behaviors of *flp-3* [38].

## Materials and Methods

### *C. elegans* Strains

*C. elegans* strains were maintained on nematode-growth media (NGM: 51 mM NaCl, 10 mM peptone, 51 mM agar, 1 mM CaCl_2_, 1 mM MgSO_4_, 25 mM KPO_4_ (pH 6.0), 13 µM Cholesterol) plates seeded with *E. coli* OP50 at 15 – 20°C, according to standard procedures [39]. Strains used are listed in **Table S1**. The strains selected from previous publications have validated the associated neuropeptide behavior. For *pdf-1* associated assays, the strain UR954 (*pdf-1(tm1996)*III; *him-5(e1490)*V) creates a *pdf-1* loss of function (lof) behavior by a 588 base pair gene deletion resulting in a splice donor variant. The control strain for *pdf-1* behaviors was CB4088 *him-5(e1490)*V, which creates a higher incidence of males by a point mutation (G to A) resulting in a splice acceptor variant. *flp-3 lof* behavior was tested by JSR99 (*flp-3(pk361)*X; *him-8(e1489)*IV) whose *flp-3* allele results in a 439 base pair deletion resulting in a suspected frame shift. The *him-8* allele CB4088 (*him-8(e1489)*IV) selected results in a higher incidence of males by a point mutation (G to A) resulting in a missense mutation. The Bristol wildtype isolate (N2) was used as the control for the INS behavior tests since they were based in hermaphrodite behavior. *ins-6 lof* behavior was tested by IV302 (*ins-6(tm2416)*II; kyEx2595[*str-2*p::GCaMP2.2b *+ unc-122*p::GFP]), where the tm2416 allele results in a 316 base pair deletion that results in a splice donor variant. *ins-22 lof* behavior was tested by RB2594 (*ins-22(ok3616)*III) which creates a ∼700 base pair gene deletion resulting in a putative frame shift.

JSR188 was generated by mating UR954 and UR930 animals. UR930 (*pdfr-1(ok3425)*III; *him-5(e1490)*V) results in *pdfr-1* loss of function by a 605 base pair deletion resulting in a splice acceptor variant. The cross was monitored using validated primers from Wormbase.org; *pdf-1(tm1996)*external left (CCGTGGAGGTACCAGGTTAA), *pdf-1(tm1996)* external right (TCAAGGTCTCGCCTCTAGGA), *pdfr-1(ok3425)*external left (CGTGGAATCATCGCTACCTT), and *pdfr-1(ok3425)* external right (TTTATGCAGGCTTATTGCCC). The final generated strain was confirmed by sequencing using the primers *pdf-1(tm1996)* internal left (CCATTCCGATGAGTCCGTTG) and *pdfr-1(ok3425)* internal left (TTTACTCCTTGACGGGAACG). To test the FLP-3 receptor FRPR-16 associated behavior, JSR152 (*flp-3(pk361)*X; *frpr-16(gk5305*[*lox*p *+ myo-2*p::GFP::*unc-54 3’ UTR + rps-27*p::neoR::*unc-54 3’ UTR + lox*p]II)) was used, creating a *frpr-16 lof* mutation by deleting 1685 base pairs, resulting in a splice acceptor variant. JSR166 was generated by mating IV302 and HC196 animals. HC196 (*sid-1(qt9)*V) results in *sid-1* loss of function single base pair mutation C to T resulting in the gain of a stop codon. The cross was monitored using primers external left (TCCTTCGCCTGGAATTAGAT), external right (AGCAAAATCCGTACATTCGC) and sequenced with an internal left primer (AAACTTCCGACGGTTTATCC).

### Neuropeptide Plasmid Design and Feeding

#### Peptide Plasmid Design and Generation

DNA sequences encoding individual peptides were identified via www.wormbase.org. The endogenous processing patterns of the peptides selected are mapped in (**Figure S1**). Sequences were flanked with the endogenous cleavage sites for the EGL-3 processing enzyme, which cleaves dibasic resides. Sequences encoding MRFGKR and KRK-STOP codons were therefore placed prior to, and following the peptide codon sequences, respectively. This construct lacks the signal peptide sequence found in endogenous neuropeptides. Finally, Gateway Cloning sites attB1 and attB2 sites were attached to the ends of the sequences. These final sequences (comprised of attB1::MRFGKR::peptide::KRK-STOP::attB2) were ordered from Integrated DNA Technologies (IDT) using their DNA Oligo and Ultramer DNA Oligo services, depending on the size of the oligo ordered. Both forward and reverse sequences were ordered (**Table S2**).

Lyophilized oligos were prepared following IDT Annealing Oligonucleotides Protocol (https://www.idtdna.com/pages/education/decoded/article/annealing-oligonucleotides). In short, they were resuspended in Duplex Buffer (100 mM Potassium Acetate; 30 mM HEPES, pH 7.5; available from IDT), preheated to 94 °C to a final concentration of 40 μM. Complementary oligo sequences were then mixed in equimolar ratios and placed in a thermocycler at 94 °C for 2 minutes prior to a stepwise cooling to room temperature.

Annealed oligos were used to perform a BP reaction with pDONR p1-2 donor vector to generate pENTRY clones (Gateway Cloning, ThermoFisher). Entry clones were then recombined with pDEST-527 (a gift from Dominic Esposito (Addgene plasmid # 11518)) in LR reactions generating expression clones. The control (SCRAMBLE) was generated in an identical manner, with the sequence between the cleavage sites being amplified from pL4440 (provided by Victor Ambros, University of Massachusetts Medical School, MA) to produce a peptide that lacks sequence homology to any known peptide (**Table S2**). Purified neuropeptide expression vectors were stored at -20 °C or -80 °C for long-term storage.

For the *ins-22* experiments, the DNA coding sequence for the peptide was identified as described above, without the Gateway Cloning sites. These sequences were codon-optimized and cloned in pET-21a(+) cloning vector via XhoI/XhoI strategy by GenScript (www.genscript.com). The SCRAMBLE control sequence and plasmid were generated using the same cloning strategy by GenScript. Plasmids arrived lyophilized and were re-suspended in nuclease-free H_2_O to a concentration of 100 ng/μL and stored at -20 °C.

The plasmid maps displayed in **Figure S2a,b** were rendered by SnapGene® software (from Dotmatics; available at snapgene.com).

#### Peptide Feeding Bacterial Preparation

The neuropeptide expression vectors were transformed into competent DH5α cells (NEB® [New England Biolabs] 5-alpha Competent *E. coli* (High Efficiency) Cat# C2987I). Peptide-expressing cultures on LB agar plates (10 g/L NaCl, 10 g/L Tryptone, 5 g/L Yeast Extract, 5 g/L Agar) containing 100 µg/mL ampicillin (AMP), were stored at 4 °C for up to 2 months. From this plate, single colonies were selected for growth in 5 mL of LB media (10 g/L NaCl, 10 g/L Tryptone, 5 g/L Yeast Extract) containing ampicillin at 37 °C for 16 hours. The optical density of the grown cultures (OD_600_) was adjusted to 1.0. 75 μL of 1.0 OD_600_ peptide-expressing culture was plated onto 60 mm NGM agar plates containing 100 μg/mL AMP and 1 mM isopropyl-β-D-thiogalactoside (IPTG) at room temperature for at least 8 hours prior to use, for no longer than 1 week.

Competent iOP50 (*E. coli*, rnc14::(delta)Tn10, lacz(gamma)A::T7pol camFRT) obtained from *Caenorhabditis* Genetics Center was created using previously described methods [40]. Briefly, iOP50 stocks were streaked out onto plates containing 50 µg/mL tetracycline and LB and grown at 37°C overnight. Then, single colonies of iOP50 were isolated and used to inoculate 2.0 mL LB containing 50 µg/mL tetracycline and placed in a 37°C shaker overnight. 0.5 mL of each culture was used to inoculate 3.0 mL of LB medium without antibiotics and was grown for 2 hours at 37°C in a shaker. Each culture was then aliquoted in 0.75 mL increments into four 1.5 mL tubes and cooled on ice for 10 minutes. Next, the cells were harvested by centrifugation at 6,000 rpm for 5 minutes at 4°C. The resulting supernatant was discarded, and the cells were placed back on ice and resuspended in 1.0 mL 100 mM ice-cold CaCl_2_ solution. Cells were kept on ice for 20 minutes and then harvested by centrifugation as in the previous step. After discarding the supernatant, the cells were resuspended in 150 µL of ice-cold CaCl_2_ solution and placed on ice for immediate use in transformation.

Expression clones were transformed into competent iOP50 cultures using the New England Biolabs protocol (C2987H/C2987I). Single colonies of transformed cells were isolated and cultured overnight in 3.0 mL of LB medium containing 50 µg/mL tetracycline (TET) and 17 µg/mL chloramphenicol (CAM) at 37°C in a shaker. Cultures were frozen down as glycerol stocks and stored at -80°C. Successful transformation was verified by isolating plasmids from cultures using a Zyppy Plasmid Miniprep Kit (Zymo Research #D4019) and sending samples for whole-plasmid sequencing.

For plating the bacteria containing *PDF-1* and *FLP-3*, LB medium containing 50 µg/mL ampicillin (AMP), 50 µg/mL tetracycline (TET), and 17 µg/mL chloramphenicol (CAM) was inoculated with iOP50 expressing either the peptide of interest or the control peptide and cultured for 16-20 hours at 37°C in a shaker. 75 µL of bacteria expressing the peptide was plated onto 60 mm NGM plates containing 50 µg/mL AMP, 50 µg/mL TET, and 17 µg/mL CAM, and 1 mM IPTG. For all peptides, seeded plates were left to dry at least 8 hours at room temperature before plating worms and were used no longer than 1 week after plating.

For *In-22,* LB containing 50 µg/mL tetracycline and 50 µg/mL carbenicillin was inoculated with iOP50 expressing either the peptide of interest or the control peptide and cultured for 18-20 hours at 37°C in a shaker. 100 mm NGM plates containing 50 µg/mL tetracycline, 50 µg/mL carbenicillin, and 1 mM IPTG were plated with 1.0 mL of the corresponding culture. Seeded plates were left to dry for at least 24 hours at room temperature and were never stored for longer than 72 hours before use. Approximately one hour before transferring worms to peptide or control plates (SCRAMBLE), 200 µL of 0.1 M IPTG was added to each plate and left to dry.

#### Peptide Feeding of Bacterial Cells or Supernatant Alone

To test if the bacterial cells expressing the peptide construct or the supernatant that the cells were grown in was sufficient to restore neuropeptide-associated behavior, the cells expressing the peptide were separated from the media in which they grew and might have expressed macromolecules into. 5 mL of LB medium containing 100 µg/mL ampicillin (AMP) with DH5α cells expressing the control SCRAMBLE construct or FLP-3-9 was grown for 16 hours at 37 °C with shaking. At the same time, 5 mL of LB containing OP50 *E. coli that did* not express the peptide was grown for 16 hours at 37 °C with shaking. Afterwards, the cultures were centrifuged at 3900 RPM (rotations per minute) or 11,000 x G for 5 minutes at 4 °C to separate the cells from the media. The supernatant from the DH5α *E. coli* cells expressing the peptide was mixed with the OP50 cells not expressing the peptide, and the DH5α *E. coli* cells expressing the peptide were mixed with the OP50 cell supernatant. The cultures were adjusted to have an OD_600_ = 1.0. 75 μL of each culture (SCRAMBLE DH5α cells, SCRAMBLE DH5α supernatant, FLP-3-9 DH5α cells, and FLP-3-9 DH5α supernatant) was plated onto 60 mm NGM agar plates containing 100 μg/mL AMP and 1 mM IPTG. The plates were stored at room temperature, 20-22 °C, for at least 8 hours prior to use, for no longer than 1 week.

#### Peptide Feeding

All animals were maintained on bacterial lawns of OP50 *E. coli*, at 20 °C, on NGM agar plates until the start of experiments. Mutant or control animals were transferred onto lawns of DH5α or iOP50 *E. coli*. Animals were reared on the peptide-expressing lawns for at least 48 hours at 20 °C before assaying for rescue of mutant phenotype at the appropriate developmental stage. See each section of the behavioral assay methods for the exact peptide feeding parameters.

### Behavioral Assays

#### Excursion Assay (pdf-1)

The mate searching behavior of *pdf-1* mutants in a *him-5* background was quantified in a food leaving assay. *him-5* animals were used as wildtype control to ensure the presence of males. *him-5* and *pdf-1* animals were fed bacteria expressing no peptide (OP50), control peptide (SCRAMBLE), PDF-1A peptide, PDF-1B peptide, or 1:1 PDF-1A and PDF-1B peptide (by mixing bacterial cultures expressing these peptides). Four to six larval L4 hermaphrodites were passed onto peptide-containing plates. These worms were grown at 20 °C for approximately 4 days, until the male progeny of the initial L4 worms had developed into young adults. The day prior to the assay, young adult male *pdf-1* or *him-5* worms were singled onto 60 mm plates containing lawns of either OP50 or appropriate peptide cultures.

To determine the mate searching effects of *pdf-1*, we performed a food-leaving assay as described previously [41–43], with modifications. In this assay, individuals were placed on a small food spot, and track patterns were scored at the indicated times. Assay plates were prepared one day prior to assaying on 100 mm plates with 10 mL of Leaving Assay media (25 mM KPO_4_ (pH 6.0), 1 mM MgSO_4_, 1 mM CaCl_2_, 50 mM NaCl, 2.5 g/L Peptone, 17 g/L Agar). On center of plate, 7 µL of OP50 was added. In assays using iOP50 assay plates, 7 µL of iOP50 containing the appropriate peptide (ex. SCRAMBLE iOP50 assay plates made for animals fed SCRAMBLE iOP50) was added. Plates were lightly covered and stored at room temperature overnight (∼20⁰C, < 40% humidity).

On the day of the assay, a single male was plated on the assay plate and placed in a dark, 20°C incubator. At 2, 6, and 24 hours after the worms are plated, plates were scored for male worm tracks. Tracks were scored based on their distance from the center of the plate [41]. “Never left food” indicates the absence of tracks outside the food spot. “Minor excursion” indicates that the tracks never extended beyond 10 mm from the food. “Major excursion” indicates the presence of tracks past the 10 mm boundary. We quantified the food leaving behavior at three different time points (2, 6, and 24 hours) [42].

We additionally quantified the mate searching defect of *pdf-1* as a Probability of Leaving per hour (P_L_/hr) [43]. In this method, we reduce the analysis of the leaving behavior to a single value, by scoring the worms leaving the food lawn as Leavers (traveling beyond 35 mm from the food source) or Non-Leavers (did not travel further than 35 mm from center of food). Worm in the > 35 mm from the center of food bin were scored as “Leavers” [43]. The probability of leaving per hour (Probability of Leaving (P_L_)) was calculated using an R script developed in [42], first cited in [43], with assistance by (Personal Communications to Barrios, 2020). P_L_ was calculated by the “hazard obtained by fitting an exponential parametric survival model to the censored data using maximum likelihood” [43].

#### Avoidance Assay (flp-3)

Avoidance drop assays were performed as described previously [38, 44]. This was performed on *him-8* or *flp-3;him-8* loss of function male animals. Fifty to sixty L4 male worms were picked by sex to isolate the males and stored at 20 °C for at least 5 hours to overnight to be assayed as young adults. One to four hours prior to the assay, the lids of unseeded plates were tilted to allow any excess moisture to evaporate off the plates. At the time of the assay, 10 or more males were transferred onto each of the dried, unseeded NGM plates. A drop of either water or 1 µM ascr#8 (a nonvolatile *C. elegans* mating pheromone) was placed on the tail of forward moving animals, and their response was scored as either an avoidance response, or no response. The avoidance index was calculated as:

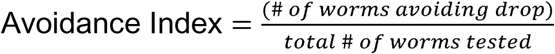

#### Chemotaxis Assay (ins-6)

A salt chemotaxis assay was performed as previously described in [45],[46], to test the functionality of *ins-6* after rescue-by-feeding. Wildtype (N2 Bristol) and *ins-6* loss of function worms were fed either a control peptide (SCRAMBLE) or INS-6 peptide. This was done throughout the animals’ lives by passing 4-6 L4 larval onto an NGM plate containing 100 μg/μL AMP and 1 mM IPTG with 75 µL of peptide OD_600_ = 1.0. These worms were grown at 20 °C for about 4 days until the progeny of the initial L4 worms were young adults.

Chemotaxis plates were prepared the day prior to testing, by adding 10 mL of agar (5 mM KPO_4_ (pH 6.0), 1 µM MgSO_4_, 1 µM CaCl_2_, 20 g/L agar, and 8 g/L Difco Nutrient Broth) to 100 mm petri dishes. To prepare the “high salt” plates, 750 mM NaCl was added prior to plate pouring. The high salt plate were stored at 4°C for up to one month, while the Chemotaxis Plates were made one day prior to testing. To create the high and low salt plugs, the back end of a Pasteur pipette was used to punch a 5 mm plug out of each plate. These 2 plugs were placed on opposite edges of the plate. A 10 mm radius was be drawn around the location of the plugs. Plates were stored, lightly covered, overnight at room temperature (∼20°C, < 40% humidity).

On the day of the assay, young adult worms, either control worms fed OP50 *E. coli* or peptide-fed worms, were washed off peptide plates with M9. Worms were allowed to settle by gravity for 4 minutes before removing the supernatant, washing the worms. Worms were then washed with Chemotaxis Buffer (5 mM KPO_4_ (pH 6.0), 1 mM MgSO_4_, 1 mM CaCl_2_) three times.

To prepare the Chemotaxis plates for the assay, high and low salt plugs were removed with forceps, and 1 µL of 0.5 M sodium azide (NaN_3_) was added to the plage where the plugs had rested. Approximately 30 µL of worms in Chemotaxis Buffer were spotted onto the edge of the agar, between the location of the high and low salt gradient. Worms were left to chemotax for 1 hour before scoring the number of worms within the 10 mm radii of high and low salt areas using the below choice index to assess the behavior of animals that had made a choice between the two salt location:

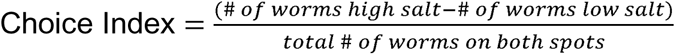

#### Olfactory Chemotaxis Assay (ins-22)

Chemotaxis assays were performed using worms that were transferred at the L4 stage to plates seeded with iOP50 containing a plasmid expressing either a control sequence (SCRAMBLE) or the peptide of interest (INS-22). These experiments were performed on wildtype (N2 Bristol) animals or *ins-22* loss of function animals. After 48 hours of exposure, we tested the naïve chemosensory preference of Day 2 Adults using a protocol based on previously published assays [44]. Briefly, assays were performed on unseeded 100mm NGMs. On the back of each plate, two marks were made on opposite sides of the plate, approximately 5mm from the edge. 1 μL of sodium azide (Thermo Fisher) was placed on both spots and allowed to dry before adding 1 μL of 0.1% or 10% butanone (Sigma Aldrich) diluted in ethanol on one mark and ethanol on the other. Using M9 buffer, worms were washed off their plates and subsequently washed three times to eliminate any leftover bacteria that could impact baseline behavior. Then, worms were placed near the bottom center of the plate, equidistant between the two marks, and allowed to chemotax for an hour. Chemotaxis indices were calculated as follows:

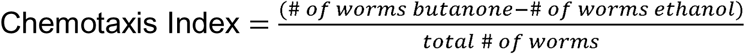

### Statistical Analysis

#### Power Analysis

Prior to the study, a post-hoc power analysis (ClinCalc LLC., Chicago, IL, USA) was performed using preliminary data from OP50 *E. coli* fed worms to determine the number of samples required to establish a difference between the wildtype controls and the neuropeptide loss of function mutants. This was performed using a 0.05 alpha rate.

For the excursion assay comparing *pdf-1 lof* animals in a *him-5* background compared to *him-5* males, a dichotomous power analysis was used to compare non-leavers (< 35 mm) to leavers (> 35 mm). n = 39 (where n = 1 is one male on one assay plate) was required for 100% powered experiment and 13 for power > 80%. A sample size of n = 50 samples per genotype and feeding condition was selected.

For avoidance behavior towards ascr#8 comparing wildtype control *him-8* animals to *flp-3 lof* males in a *him-8* background, n = 3 samples (n = 1 is one plate of ten worms) was appropriate to have a 100% powered test, using a continuous power test. A sample size of N = 10 was selected. Points plotted on graphs represent n = 1.

For the salt chemotaxis assay with *ins-6 lof* animals, n = 4 samples (n = 1 is one plate of 100-200 worms) were required for power > 80% and 13 for 100% power. A sample size of n = 10 was selected. Points plotted on graphs represent n = 1.

For chemotaxis behavior of *ins-22* animals, the behavior of SCRAMBLE fed iOP50 animals towards 10% butanone was used to determine that 20 samples (n = 1 is one plate of 100-200 worms) were required for 97.5% power and 11 samples were required for power > 80%. A sample size of n > 15 was selected. Points plotted on graphs represent n = 1.

#### Reproducibility

In all experiments, the investigators were blinded to genotype. Young adult animals were tested in all behavioral assays. For the behavioral tests of *flp-3* and *ins-6*, the assay was repeated over at least 3 days, with no more than 3 replicates in a day, until at least 10 plates had been tested. For *ins-22*, the assays included testing 5 plates per day, for a total of three biological replicates (n = 15). For *pdf-1, t*he assay was performed across at least two days per condition, with at least n = 50, wherein each assay plate with one male is equal to n = 1. When possible, the results of an individual n are displayed on the graphs.

#### Statistical Comparison

Statistical analysis was performed in GraphPad Prism (version 10.4.1). Prior to any statistical comparison, the normality and frequency distribution were assessed for each of the groups. If these tests showed normal distributions, a Student’s unpaired two-tailed t-test followed by Bonferroni’s correction was applied; this was used for *ins-6* and *ins-22* results. If the data was non-parametric, as was often the case with the *flp-3* data, a Mann-Whitney test was performed followed by Bonferroni’s correction. For the *pdf-1* tests where animals were fed different types of peptides, a One-way ANOVA with Dunnett’s post-hoc test was used to compare within genotype results. For all graphs, the average was plotted with the standard error of the mean (SEM).

## Results and Discussion

### Bacterial Delivery of Neuropeptide Vectors Rescues Behavior in Loss of Function Mutants

To demonstrate the efficacy of the neuropeptide delivery by bacterial feeding, we tested the behavior of loss of function (*lof*) animals from each of the three neuropeptide classes in *C. elegans*. Each of these behavioral assays was paired with the appropriate wildtype control. For male animal behavior, *him-5* or *him-8* lines were used to produce a higher incidence of males (him) than the wildtype population of 0.1% males [47, 48]. The choice of which *him* mutant to use depended on the genetic position of neuropeptide gene being tested and other the position of other genes of interest used in the initial studies establishing the associated neuropeptide behavior.

#### Male Excursion Behavior (pdf-1)

Wildtype males display a characteristic exploratory behavior when left on a lawn of food, as well-fed males leave food in search of mates [42, 43]. The pigment dispersing factor PDF-1 (also known as NLP-74) neuropeptide plays a significant role in male mate-searching behavior, regulating neural circuits that control the choice between two predominant male interests, finding food and finding mates [42, 49]. The *pdf-1* precursor encodes two neuropeptides: PDF-1A (SNAELINGLIGMDLGKLSAVG-N terminus) and PDF-1B (SNAELINGLLSMNLNKLSGAG-N terminus) [50, 51]. The males used in this behavior were generated by a *him-5 lof* background mutation. *pdf-1 lof* males did not leave food as readily as wildtype worms, suggesting that these worms do not display exploratory behavior, as previously described (**Figure S2a,b**) [42]. When fed either DH5α *E. coli* expressing individual PDF-1 peptide (or a 1:1 combination of both), *pdf-1 lof* had a higher P_L_ compared to SCRAMBLE fed *pdf-1 lof* males (**Figure 1a, S2c**). However, PDF-1B and a 1:1 ratio of both peptides resulted in a significantly different P_L_ from *pdf-1* males fed PDF-1A alone. While the P_L_ of peptide-fed males were significantly higher than *pdf-1 lof* males fed SCRAMBLE, they were still significantly lower than wildtype males fed SCRAMBLE (**Figure 1a**).

**Figure 1:**
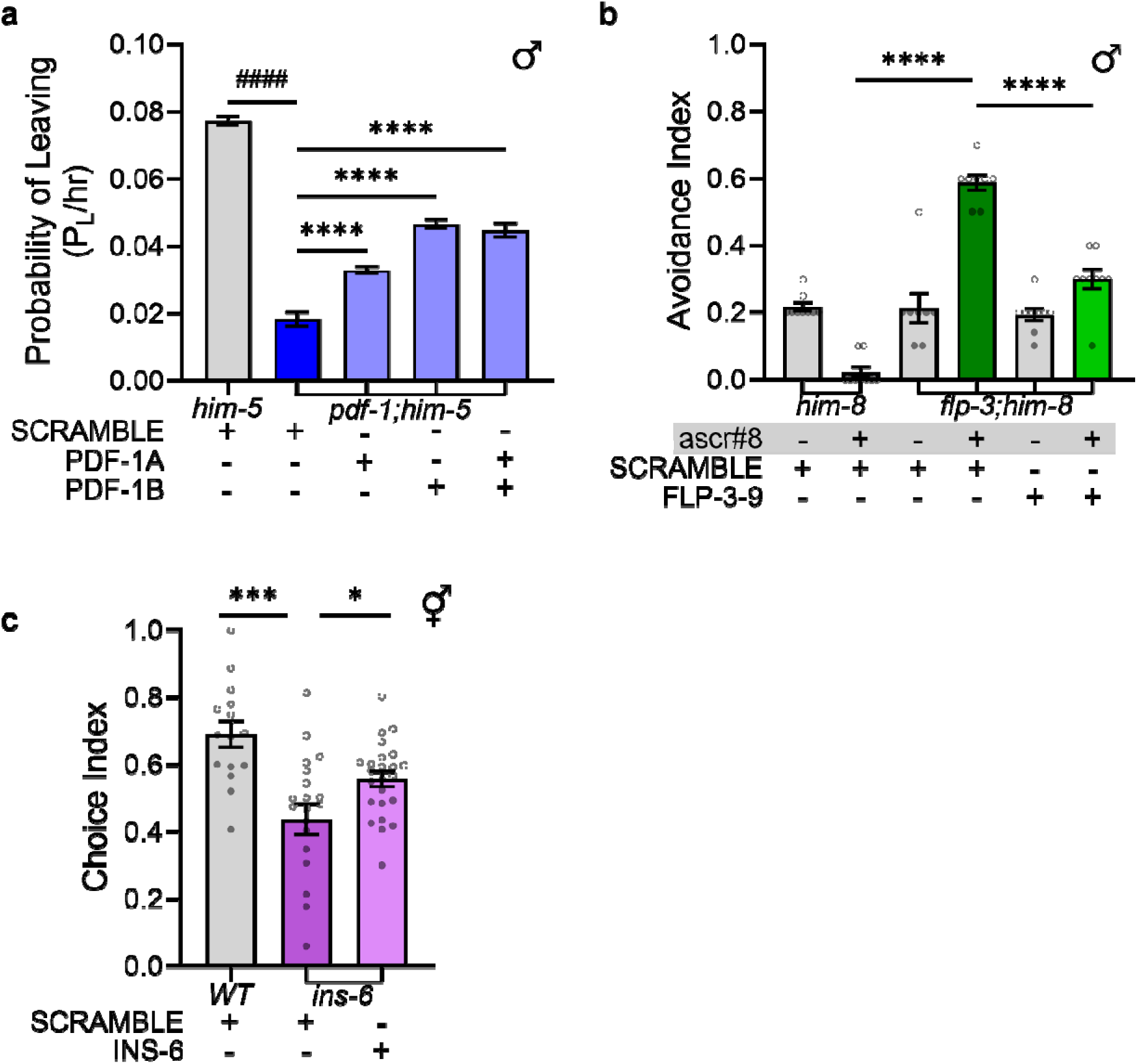
Behavioral of neuropeptide associated behavior in loss of function mutant background. **A)** *pdf-1;him-5 lof* mutant males fed a SCRAMBLE peptide have a significantly lower probability of leaving food when compared to wildtype control *him-5* males (^####^p ≤ 0.0001, unpaired t-test). When fed any of the *pdf-1* produced peptides in a DH5α *E. coli* vehicle, the probability of leaving is significantly higher than the SCRAMBLE fed *pdf-1* males (****p ≤ 0.0001, One-way ANOVA with Dunnett’s post-hoc test). **B)** When exposed to a water solvent control (ascr#8 -), both *him-8* and *flp-3* males act the same. When exposed to ascr#8 (ascr#8 +), wildtype *him-8* males do not avoid the cue while *flp-3;him-8 lof* males do. When fed FLP-3-9 peptide in a DH5α *E. coli* vehicle, the avoidance towards ascr#8 by *flp-3 lof* males is significantly decreased (****p ≤ 0.0001, Mann-Whitney test with Bonferroni correction, new p-value ****p ≤ 0.00005). **C)** Wildtype N2 animals are more attracted to high 750 mM NaCl than *ins-6 lof* animals. When fed INS-6 in a DH5α *E. coli* vehicle, the attraction of *ins-6 lof* animals to NaCl is significantly increased (*p ≤ 0.05, ***p ≤ 0.001, unpaired t-test with Bonferroni correction, new p-value *p ≤ 0.025, ***p ≤ 0.0005).

#### Pheromone Avoidance (flp-3)

*C. elegans* produce and release pheromones as a means of chemical communication to signal a variety of features to other animals, such as food or mate presence [52]. One such pheromone is ascaroside #8 (ascr#8) [53]. We have previously worked to characterize the behavior of animals towards this hermaphrodite produced pheromone and have found that wildtype hermaphrodites avoid ascr#8 while males are attracted to it [38]. We have further identified that males deficient in the neuropeptide gene *flp-3* lose their attraction towards ascr#8 and start to avoid the cue (**Figure S2d**) [38]. The males of this behavior were created in a *him-8 lof* background. The *flp-3* neuropeptide precursor creates 10 different neuropeptides, yet only 2 are involved in the male behavior towards ascr#8 [38]. Both the second and the ninth peptide (FLP-3-2 and FLP-3-9) produced by *flp-3* are involved with suppressing the basal avoidance behavior seen in *flp-3 lof* males, yet only FLP-3-9 is necessary for restoring the attraction towards ascr#8 [38, 54]. When we fed male *flp-3 lof* animals DH5α *E. coli* expressing FLP-3-9 (NPENDTPFGTMRFG-N terminus), we saw a significant suppression of avoidance behavior towards ascr#8 (**Figure 1b**).

#### Salt Chemotaxis (ins-6)

Insulin and insulin-like peptides serve signaling functions in *Drosophila* and *C. elegans* homologous to the human insulin-like growth factor (IGF), which regulates FOXO activity [55]. In *C. elegans, ins-6* encodes only one processed INS-6 peptide (VPAPGETRACGRKLISLVMAVCGDLCNPQEGKDIATECCGNQCSDDYIRSACCP-N terminus) [56, 57], though the gene was originally postulated to encode two putative proteins [1, 23]. The formation of a single longer peptide is common in the insulin-like class of peptides [58]. *ins-6* functions in dauer formation [56, 58] and sensory modulation of large fluctuations in salt concentration, with loss of *ins-6* causing dysfunction of NaCl attraction [59]. While wildtype hermaphrodite worms were attracted to high concentrations of salt (750 mM), there was a significant decrease in attraction in *ins-6* mutant animals (**Figure S2e**) [59]. Wildtype and *ins-6 lof* animals fed SCRAMBLE peptide also had significantly different levels of salt chemotaxis (**Figure 1c**). Feeding of DH5α *E. coli* expressing INS-6 peptide to *ins-6 lof* animals resulted in an in attraction to high salt concentration (**Figure 1c**).

### Supernatant from Bacteria Expressing Peptide May Be Sufficient for Behavioral Rescue

We tested whether the cells or the supernatant from a grown liquid culture expressing the neuropeptide construct was sufficient for behavioral restoration (**Figure S3**). We selected the *flp-3* associated avoidance behavior of ascr#8 to screen the different culturing conditions. We found that while the behavior of *him-5* males fed any combination of the initial culture of DH5α cells or supernatant expressing a peptide construct (SCRAMBLE or FLP-3-9) did not alter their lack of avoidance towards ascr#8. However, *flp-3;him-5 lof* males did exhibit reduced avoidance to ascr#8 when they were fed a combination of the supernatant of DH5α cells expressing the FLP-3-9 peptide with OP50 cells that did not express any peptide. These results suggest that the peptide macromolecule of interest that restores the neuropeptide associated behavior is excreted from the bacterial cells during the initial growth of the culture. This also supports that the cells do not need to be induced with IPTG (which is only present in the plates and not the liquid cultures) to express this macromolecule.

This lack of induction requirement might not entirely be surprising. Other researchers using *E. coli* as a vehicle for expressing a designed construct have found it to have leaky expression prior to induction, whether by IPTG or another induction agent [60, 61]. These studies found that while the amount of dsRNA or enzyme produced did increase after induction, the uninduced state was sufficient for the creation of these macromolecules. This would mean that the bacterial cells are able to create the expression vector macromolecules regardless of the addition of IPTG.

### Different *E. coli* food sources lead to Neuropeptide Behavior Rescue

While the previous experiments in this study have used DH5α *E. coli* expressing peptides to shape neuropeptide related behaviors, there are cases when using different *E. coli* strains may be desirable. Given the differences in nutrient composition of the different *E. coli* strains and worms preferring more nutritious bacteria [62], expressing peptides in HB101 or HB115 strains offers viable alternatives. To most closely compare to other laboratory studies, the ability to use the widely used uracil deficient OP50 *E. coli* strain as the neuropeptide delivery system is desirable [39]. Recently an IPTG inducible strain of OP50 has been created, iOP50, which can be cultured to create chemically competent cells [40]. All constructs presented in this section were transformed into iOP50 before being fed to the *C. elegans*.

For testing male *pdf-1* associated leaving behavior, we first fed the animals on iOP50 peptide plates before transferring them to a leaving assay plate that is seeded with OP50 containing no peptide. Under these conditions, male *pdf-1 lof* animals fed PDF-1 peptides from iOP50 left food at a higher rate than *pdf-1 lof* animals fed a SCRAMBLE peptide from iOP50 (**Figure 2a**). However, the extent of food leaving by males after being fed iOP50 with peptide was lower than observed when animals were fed DH5α *E. coli* (**Figure 1a**). This may be because animals are tested over the course of 24 hours and the level of peptide in their bodies may decrease over time. To address this, we tested animals fed iOP50 with peptide and then transferred them to iOP50 assay plates seeded with the same peptide they were previously fed (**Figure 2b**). By doing this, the trend of *pdf-1 lof* animals fed PDF-1 leaving food at a higher rate continued, and the overall rates of leaving food also increased (**Figure 2b**).

**Figure 2:**
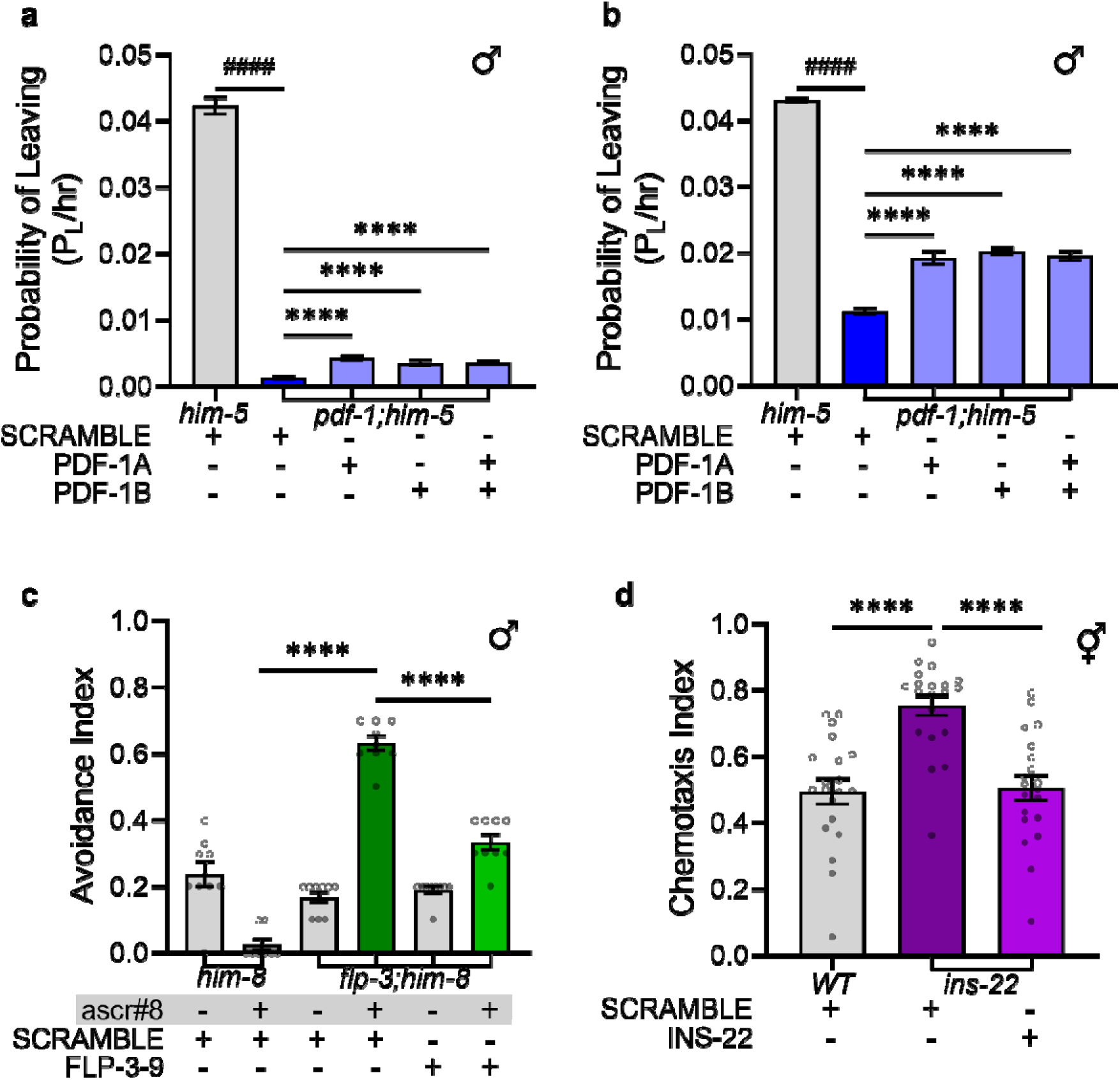
Neuropeptide rescue by feeding can be accomplished in different *E. coli* vehicles. **A)** *him-5* males fed a SCRAMBLE peptide in iOP50 *E. coli* leave food at a higher rate than *pdf-1;him-5 lof* males (^####^p ≤ 0.0001, unpaired t-test). *pdf-1;him-5 m*ales fed PDF-1 peptides in iOP50 and then plated on an assay plate containing OP50 *E. coli* without peptide, do leave food at a higher rate than *pdf-1* SCRAMBLE iOP50 *E. coli* fed animals (****p ≤ 0.0001, One-way ANOVA with Dunnett’s post-hoc test). **B)** *him-5* males fed SCRAMBLE peptide in iOP50 *E. coli* and then placed on an assay plate seeded with SCRAMBLE peptide in iOP50 *E. coli* leave food at a higher rate than *pdf-1;him-5 lof* males (^####^p ≤ 0.0001, unpaired t-test). *pdf-1;him-5* males fed PDF-1 peptides in iOP50 and then plated on an assay plate containing iOP50 *E. coli* expressing the corresponding peptide, leave food at a higher rate than *pdf-1;him-5 lof* animals fed SCRAMBLE iOP50 *E. coli* and at a higher rate than animals on assay plates without peptide (****p ≤ 0.0001, One-way ANOVA with Dunnett’s post-hoc test). **C)** When exposed to a water solvent control (ascr#8 -), both *him-8* and *flp-3;him-8* males act the same. When exposed to ascr#8 (ascr#8 +), wildtype *him-8* males do not avoid the cue while *flp-3 lof* males do. When fed FLP-3-9 peptide in iOP50 *E. coli*, the avoidance towards ascr#8 by *flp-3 lof* males is significantly decreased (****p ≤ 0.0001, Mann-Whitney test with Bonferroni correction, new p-value ****p ≤ 0.00005). **D)** Animals deficient in *ins-22* show significantly stronger chemotaxis towards 10% butanone than *ins-22 lof* animals fed INS-22. The *ins-22 lof* animals fed INS-22 have a chemotaxis index no different than wildtype animals (^ns^p ≥ 0.05, ***p ≤ 0.001, unpaired t-test wildtype SCRAMBLE to INS-22, Mann-Whitney test comparisons to *ins-22* fed SCRAMBLE with Bonferroni correction (new p-value ^ns^p ≥ 0.025, ****p ≤ 0.00005).

*flp-3 lof* males exposed to ascr#8 after being fed FLP-3-9 in iOP50 lacked avoidance to the cue while males fed the SCRAMBLE peptide continued to avoid it (**Figure 2c**). These results matched the observed behavior of animals fed FLP-3-9 in DH5α *E. coli* (**Figure 1b**). For *flp-3*, males are transferred directly from iOP50 peptide plate onto an unseeded NGM assay plate for no more than 30 minutes during the testing. Due to this, it was not necessary to reapply the iOP50 peptide during testing.

We also tested the delivery of peptide using a different neuropeptide expression vector. To test the behavior associated with INS-22, a new backbone vector of pET-21 was procured to circumvent the use of Gateway Cloning and the pDest-527 vector (**Figure S4a**). In this new neuropeptide expression vector, the peptide endogenous peptide sequence flanked by the peptide cut sites was ordered as a complete plasmid in a pET-21 vector (**Figure S2b**). This vector was used to test the behavior of INS-22 (MHTTTILICFFIFLVQVSTMDAHTDKYVRTLCGKTAIRNIANLCPPKPEMKGICSTGEYP SITEYCSM-N terminus). Unlike INS-6, INS-22 antagonizes the DAF-2 neuropeptide receptor [63, 64]. We discovered that animals lacking *ins-22* exhibit increased naive chemotaxis towards a neutral concentration of 10% butanone compared to wildtype animals (**Figure S4c**). This behavior persists in animals fed the SCRAMBLE peptide in iOP50 *E. coli* (**Figure 2d**). When fed the INS-22 peptide, the *ins-22 lof* animals displayed naive chemotaxis similar to wildtype animals, indicating that peptide feeding rescued normal naive chemotaxis towards 10% butanone (**Figure 2d**).

### Neuropeptide Rescue by Feeding Requires the Cognate Neuropeptide Receptor

To ensure that the observed behavior is resulting from the neuropeptide being fed to the animal, activating the neuropeptide receptor, we tested the behavior of animals deficient in the neuropeptide receptor. Pairing neuropeptides ligands to their cognate receptors is an ongoing effort in peptidomic research. Recent efforts have uncovered many of the individual neuropeptide pairings and the range over which these connections functions [26, 65, 66]. For the current study, we were interested in studying the function of receptors that have one peptide ligand. This would eliminate the class of insulin-like peptides which bind to DAF-2 from this current study [23, 63].

PDF-1 peptides binds primarily to one neuropeptide receptor PDFR-1 [42, 67]. Though PDFR-1 also binds to peptides produced by *pdf-2* (also known as *nlp-37*), these ligands are not involved in behavior of males leaving food to search for mates [42, 66]. *pdfr-1 lof* mutants in a *him-5* background do not leave food to search for mates (**Figure S5a,b**), even when fed excess peptide (**Figure 3a, S5c**). The double mutant males of the *pdfr-1* and *pdf-1* neuropeptide, partially left food to search for mates when fed PDF-1A but not when fed the other peptides (**Figure 3a**). Though PDF-1A was able to promote the male leaving behavior, we do not believe this was to a biologically significant level since the rates of leaving food were lower than the rate than *pdf-1 lof* males left food. This probability of leaving food of *pdf-1;pdfr-1* males was not to the rate of *pdf-1* males leaving food when fed a SCRAMBLE peptide (**Figure 3a**). The absence of food leaving to search for mates, even when receptor mutants are fed the neuropeptide, suggests that there are no off-target effects of the neuropeptide impacting other receptors.

**Figure 3:**
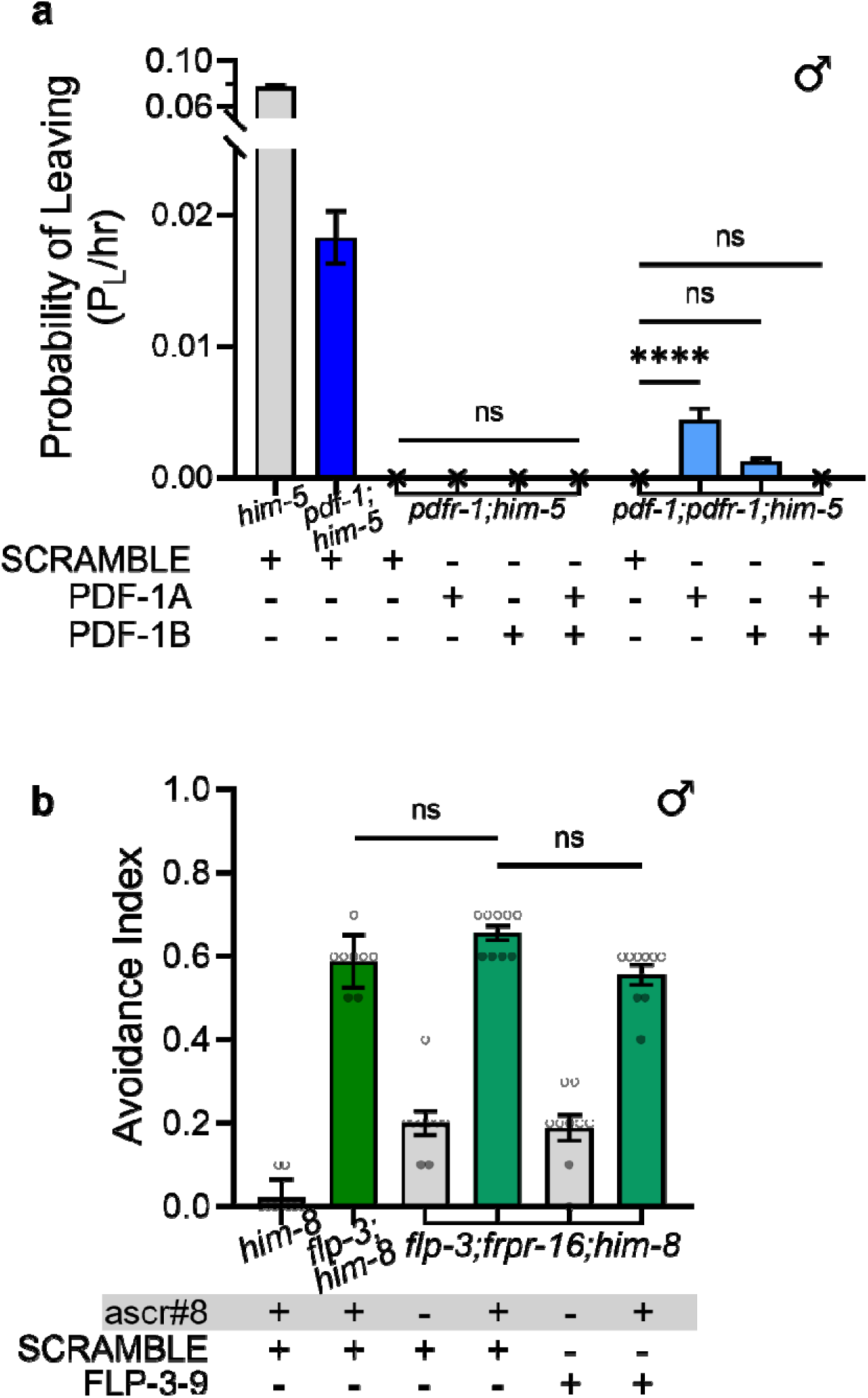
Neuropeptide receptor required for neuropeptide feeding. **A)** *pdfr-1;him-5 lof* animals do not leave food more than 35mm, even if fed PDF-1 peptides in DH5α *E. coli*. Double mutants of *pdf-1;pdfr-1* in a *him-5* background also did not leave food at the rate of *pdf-1* animals fed SCRAMBLE, though *pdf-1;pdfr-1;him-5* animals fed PDF-1A DH5α *E. coli* do have a higher incidence of leaving food than SCRAMBLE fed animals (^ns^p ≥ 0.05, ****p ≤ 0.0001, One-way ANOVA within genotypes with Dunnett’s post hoc test). **B)** Double mutant *flp-3;frpr-16* males in a *him-8* background fed bacteria expressing FLP-3-9 avoid ascr#8 to the same level as SCRAMBLE DH5α *E. coli* fed animals (^ns^p ≥ 0.05, Mann-Whitney test with Bonferroni correction, new p-value ^ns^p ≥ 0.025).

The *flp-3* neuropeptide gene makes 10 different peptides that bind to 4 different receptors [38, 66]. Neuropeptide receptors DMSR-1-2 and DMSR-7-1 bind to many of the *flp* class of neuropeptides including *flp-3*, while FRPR-16-1 binds *flp-3* and *nlp-23* peptides and NPR-10-1 binds *flp-3* and *nlp-23* peptides [66]. We have previously identified that the receptors FRPR-16 and NPR-10 are involved in the *flp-3* male behavior towards ascr#8 [38]. The FLP-3-9 neuropeptide binds activates the FRPR-16 receptor at lower concentrations than the NPR-10 receptor [38], so we elected this as our neuropeptide receptor candidate for FLP-3-9. We have previously demonstrated that without FRPR-16 males will avoid ascr#8 [38]. When we generated a double mutant of *flp-3;fprpr-16* we observed avoidance of ascr#8 in animals fed OP50 *E. coli* without peptide (**Figure S5a**). When fed FLP-3-9 peptide in DH5α *E. coli*, the avoidance behavior is sustained (**Figure 3b**). These data shows that while FLP-3-9 binds to multiple receptors, FRPR-16 loss of function is sufficient to alter male behavior towards ascr#8.

### Behavioral Impacts of Neuropeptide Overexpression Depend on the Peptide Identity

One question that arises with this feeding method is, how much neuropeptide makes its way into the worms and what would happen if there were an overexpression of neuropeptide? Quantifying the amount of neuropeptide produced endogenously is a difficult open question in the peptidomics field. One indication of how much of a particular neuropeptide is required is by looking at the neuropeptide precursor gene. Some neuropeptides are encoded multiple times in the same gene, like *flp-6* encoding KSAYMRFG-N terminus six times [68]. One method used to detect the presence of neuropeptides is liquid chromatography mass spectrometry (LC-MS) [19, 69–71]. While this method is extremely precise for detecting known and novel neuropeptides, it lacks the ability to quantify the amount of each neuropeptide present *in vivo* (personal communication with Sven Van Bael and Lisabet Timmermann, KU Leuven). One way to test the effects of what is likely an overexpression of neuropeptide is by feeding the neuropeptide-containing DH5α bacteria to wildtype animals.

We tested the appropriate wildtype controls for each previously assessed behavior. We observed an increase in the leaving behavior of *him-5* male worms fed PDF-1B and a decrease when fed PDF-1A in DH5α *E. coli* (**Figure 4a, S6a**). This behavioral difference disappears when both peptides are fed in combination (**Figure 4a**). These data suggest that different PDF-1 peptides play varying roles in mating behavior. Next, we examined the effects of this behavior using the iOP50 *E. coli* delivery. When males were fed an excess of iOP50 peptide and subsequently placed on an iOP50 assay plate, their rates of leaving food (**Figure 4b, S6b**) did not increase. Wildtype animals fed PDF-1 peptides produced by iOP50 also did not show an increased leaving rate of food when tested on an OP50-containing assay plate (**Figure S6c,d**). In fact in both scenarios, the rates of leaving food decrease. This suggests that there may be a precise balance of neuropeptide required for proper behavioral tuning.

**Figure 4:**
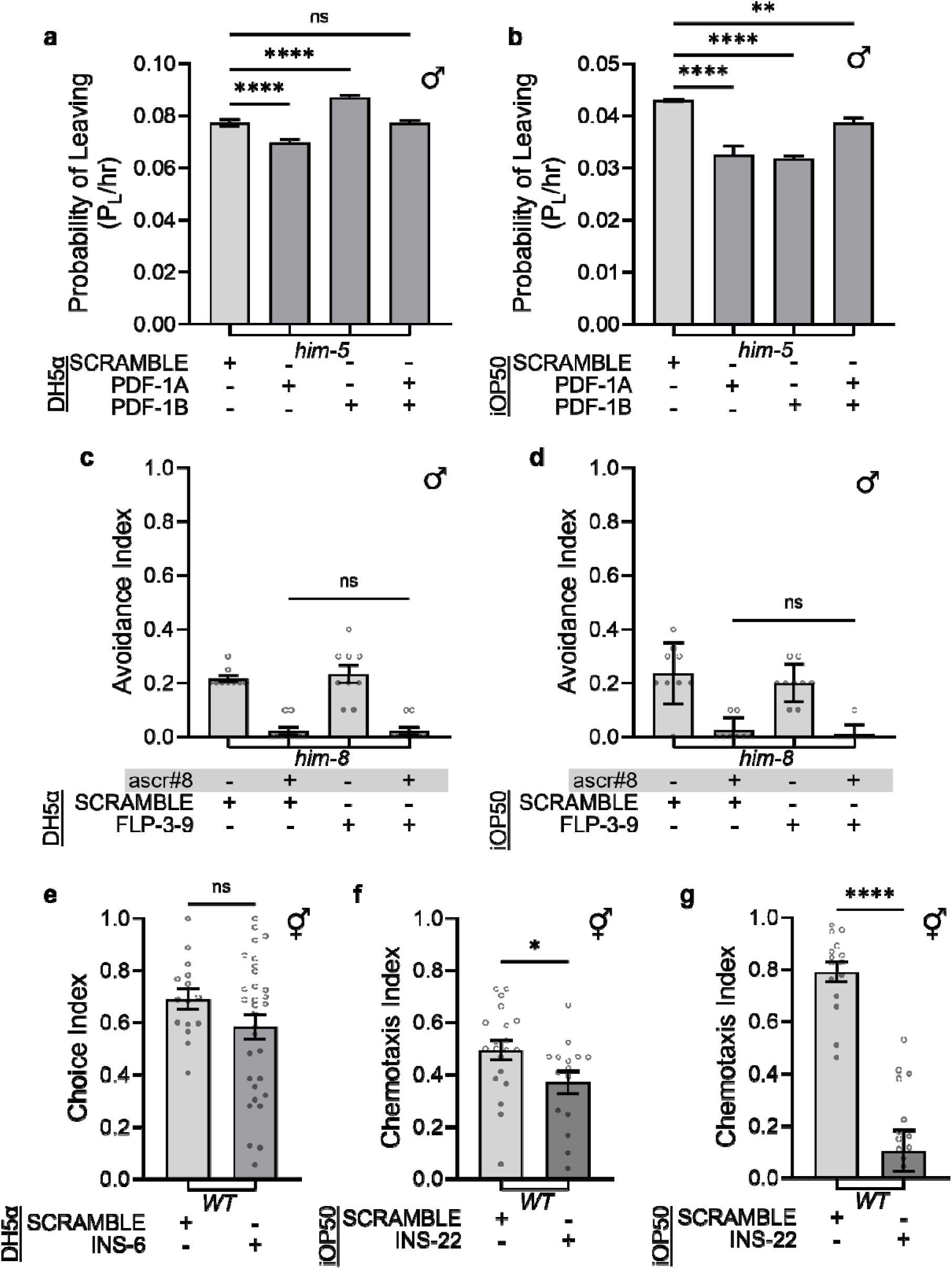
Behaviors resulting from overexpression of neuropeptides depend on the neuropeptide. **A)** *him-5* males fed PDF-1A in DH5α *E. coli* had a significant decrease in their probability of leaving food rate compared to animals fed SCRAMBLE peptide. PDF-1B *him-5* animals had a significant increase in P_L_/hr when compared to SCRAMBLE fed animals. When PDF-1A and -1B were fed to animals in equal combination, their probability of leaving food was no different from SCRAMBLE fed animals (****p ≤ 0.0001, One-way ANOVA with Dunnett’s post-hoc test). **B)** *him-5* males fed PDF-1 peptides in iOP50 *E. coli* and then tested on iOP50 assay plates containing a corresponding peptide, do not have excursions leaving food more than 35 mm at higher rates than SCRAMBLE fed males (**p ≤ 0.01,****p ≤ 0.0001, One-way ANOVA with Dunnett’s post-hoc test). **C)** *him-8* males exposed to ascr#8 had no difference in their avoidance behavior whether they were fed bacteria expressing SCRAMBLE or FLP-3-9 in DH5α *E. coli* (^ns^p ≥ 0.05, Mann-Whitney test). **D)** *him-8* males fed SCRAMBLE or FLP-3-9 in iOP50 *E. coli* avoid ascr#8 at the same rate and behave the same towards the water solvent control as the previous studies (^ns^p ≥ 0.05, Mann-Whitney test). **E)** Wildtype hermaphrodites fed SCRAMBLE or INS-6 peptide in DH5α *E. coli* displayed the same attraction towards high 750 mm NaCl (^ns^p ≥ 0.05, unpaired t-test test). **F)** Wildtype hermaphrodites fed INS-22 peptide in iOP50 *E. coli* displayed slightly less attraction towards 10% butanone than animals fed SCRAMBLE (*p ≤ 0.05, unpaired t-test). **G)** Wildtype hermaphrodites fed INS-22 peptide in iOP50 *E. coli* were not as attracted to 0.1% butanone compared to animals fed SCRAMBLE (****p ≤ 0.0001, unpaired t-test).

When *him-8* male animals were fed FLP-3-9, their lack of ascr#8 avoidance was not changed compared to animals fed SCRAMBLE (**Figure 4c**). Together these data indicate that there are no obvious behavioral deficits across all neuropeptides being delivered in excess. Wildtype males fed FLP-3-9 iOP50 *E. coli* continued their non-avoidance of ascr#8 (**Figure 4d**).

We then tested different insulin ligands for their overexpression phenotypes. When wildtype N2 animals was fed INS-6 in DH5α *E. coli*, there was no difference in the choice index of high salt taxis (**Figure 4e**). These data suggest that INS-6 overexpression by microbial feeding does not impact the behavior of animals towards high NaCl. There has been evidence that overexpression of some insulin like peptides may impact other behaviors like dauer formation [55].

When wildtype animals were fed an excess of INS-22 iOP50 *E. coli*, their behavior towards 10% butanone was decreased compared to animals fed SCRAMBLE (**Figure 4f**). When exposed to a highly attractive concentration of 0.1% butanone, wildtype animals fed INS-22 in iOP50 *E. coli* significantly reduced their attraction towards the cue (**Figure 4g**). While differences in behavior towards varying concentrations of butanone have been previously established [72, 73], the impact of INS-22 on these behaviors has not been implicated in detail. Differences between the overexpression effects of the *ins* peptides may be due to the different interactions with the DAF-2 receptor, INS-6 is a strong agonist and INS-22 is an antagonist [63, 64].

### Neuropeptide Feeding Does Not Utilize RNAi-based Mechanisms

Our method of neuropeptide study is most similar to RNA interference approaches, leading us to hypothesize that the microbes could be delivering RNA to the *C. elegans*. RNAi functions to knock down a gene by delivering dsRNA that interferes with the endogenous mRNA of that gene [74–76]. RNAi was first performed in *C. elegans* [75, 76] and has since been accomplished by feeding, soaking, and injecting dsRNA [77]. The RNAi vector contains two T7 promoters going in opposite directions to create dsRNA [74, 78], while our vector contains only 1, indicating that single stranded RNA (ssRNA) should be produced. The RNAi vector is then contained in an *E. coli* strain that has been modified to make more dsRNA, such as HT115 [79, 80]. RNA produced by bacteria in the gut microbiota can be transported across the intestinal epithelium, as identified by RNAi feeding [80]. This transport has also been seen with non-coding RNA being transmitted from pathogenic bacteria to worms [81]. We hypothesize that unlike RNAi, this method can be used to introduce larger neuropeptides encoded by over 30 base pairs [82].

To assess if our neuropeptide delivery system utilizes previously identified RNAi mechanisms, we tested mutants deficient in the ability to transport RNA. We specifically looked into *sid-1* (systemic RNA interference defective protein 1) loss of function mutants. SID-1 selectively binds to dsRNA of varying (selecting towards longer) length and not ssRNA or dsDNA [83–86]. We expect that our rescue by feeding approach should not be effected by the loss of SID-1 since we predict that our method is either relying on dsDNA, due to the selection of DH5α *E. coli* not containing T7 polymerase [87]. We found that mutants lacking *sid-1* behave like wild-type animals when receiving INS-6 peptide through feeding (**Figure S7a**). Additionally, we discovered that *sid-1;ins-6* double mutants fed the SCRAMBLE peptide display the *ins-6 lof* phenotype (**Figure S7b**). When fed INS-6, *sid-1;ins-6 lof* animals restored their NaCl attraction to in a pattern similar to single mutant *ins-6* animals, suggesting that SID-1 is not involved in our neuropeptide rescue by feeding method.

## Conclusions

We present a novel method of neuropeptide rescue that utilizes bacterial expression to deliver individual peptides for behavioral rescue phenotypes. Our technology is advantageous over transgenic studies, as it allows for functional characterizing individual peptides. This tool readily allows for testing newly de-orphanized neuropeptide/neuropeptide receptor pairings [66]. This genetic tool is built off the principles of RNAi feeding techniques, though for a restoration rather than a reduction of function, to supply worms with the peptide of interest through their food source as previously shown for the scorpion venom peptide mBmKTX to alter lifespan and egg-laying behavior in *C. elegans* [34]. Based on our results in the present study, we believe that this method can be used to restore neuropeptide function within the different class of neuropeptides in *C. elegans*.

Our future investigation will be focused primarily on uncovering the mechanisms by which the bacteria create and deliver the neuropeptide macromolecule to *C. elegans*. We believe it unlikely that the *C. elegans* are processing the DNA. Even though DH5α *E. coli* is primarily used to make DNA, and does not contain T7 polymerase, it does contain *E. coli* RNA polymerase which could recognize additional promoter sites [88]. If the *E. coli* isn’t producing the RNA, then that requires the *C. elegans* to be doing so. Like other eukaryotes, *C. elegans* have 3 types of RNA polymerases which each produce a distinct type of RNA all within the nucleus [89, 90]. For this rescue by feeding paradigm, we would expect RNA polymerase II to be producing the RNA since we expect ssRNA acting as mRNA to be produced and then fabricated into neuropeptides at the ribosome should *C. elegans* be the ones required to make the protein [90]. This would again require the neuropeptide vector to contain promoter sites to be recognized by RNA polymerase II [91–93]. If the *E. coli* is producing ssRNA, there are mechanisms by which this macromolecule can translocate into the nucleus, as it is becoming more open to investigation with the rise in mRNA vaccines [94].

Previous studies have shown that RNA produced by bacteria in the gut microbiota can be transported across the intestinal epithelium [80, 81], and our vector for neuropeptide delivery is modeled after RNAi feeding [38, 77, 78, 80]. Delivery of exogenous scorpion venom peptide proved effective in changing *C. elegans* behavior [34]. In humans, neuropeptides and neurotransmitters are produced by the gut microbiome and transferred to the nervous system for use [95]. Our neuropeptide delivery method may already make use of some mechanisms by which these other techniques function or could be using novel mechanisms of delivery. Furthermore, previous studies have shown that RNA uptake can alter transgenerational inheritance [96], despite degradation rates. Our rescue of the exploratory behavior of *pdf-1 lof* males even in the 24-hour timescale suggests that our feeding paradigm exploits similar mechanisms, allowing peptide-encoding mRNAs to remain present throughout the assay. We propose that if peptides were taken up, their degradation rates would likely impede rescue efficiency at later timepoints, unlike what we observe in our experiments, suggesting that the mechanism is similar to dsRNA feeding, with mRNA uptake driving rescue rather than peptide uptake.

Based on our results, we propose that the neuropeptide rescue-by-feeding strategy delivers mRNA that is ready for translation by the *C. elegans* cellular machinery, rather than supplying *C. elegans* with fully translated and processed peptides. The plasma membrane of a cell is an intricate complex of multiple lipid and protein molecules. Small molecules with moderate polarity diffuse through the cell membrane passively, but most metabolites and short peptides require specialized membrane transporters for translocation [97]. Given the large number of neuropeptides encoded in the genome of *C. elegans* [1], having specialized transporters for peptide transport is not feasible [98]. The most substantial evidence for this statement is that the processed INS-6 peptide is 54 amino acids in length and INS-22 is 68: the *C. elegans* intestine expresses peptide transporters that only uptake smaller, inactive di- and tri-amino acid peptide chains [98]. Thus, rescue of chemotaxis behavior by INS-6 peptide feeding and INS-22 suggests that the mechanism of rescue may be similar to double stranded RNA (dsRNA) uptake, rather than peptidergic transport across the intestinal membrane [99, 100]. Future investigations will see if there is a limit to the size of peptide that can be rescued.

Understanding neuropeptide function is essential from both the perspective of regulating neuronal (synaptic and non-synaptic) channels of communication and from a broader view of general neuronal functions throughout the brain. Revealing how a specific neuropeptide acts at both the cellular and subcellular receptor levels is a critical link in comprehending the role of neuromodulation in circuits. An enhanced experimental pipeline for investigating peptide function will facilitate progress toward understanding how neural circuit activity within the network is modulated by neuropeptides in its various states, resulting in flexible decision-making during behaviors.

## Data Availability

Strains (**Table S1**) and plasmids (**Table S2**) are available upon request. The authors affirm that all data necessary for confirming the conclusions of the article are present within the article, figures, and tables. The raw data or a summary containing the average, n, standard error measure, and associated statistical tests for all figures are available upon request.

## Acknowledgments

We thank Prof. Frank Schroeder, Cornell University, for providing ascr#8 and Prof. Victor Ambros for reagents. We want to thank Dr. Arantza Barrios for help in executing the leaving assay analysis in R. We extend our thanks to Dr. Sreekanth H. Chalasani and Dr. Douglas Portman for their insights and updates to the method for testing the phenotypes related to *ins-6* and *pdf-1*, respectively. pDest-527 was a gift from Dominic Esposito (Addgene plasmid # 11518; http://n2t.net/addgene:11518 ; RRID:Addgene_11518). We thank the *Caenorhabditis* Genetics Center for supplying some strains; the CGC is funded by NIH Office of Research Infrastructure Programs (P40 OD010440). The INS-22 strain was provided by the *C. elegans* Gene Knockout Project at the Oklahoma Medical Research Foundation, which was part of the International *C. elegans* Gene Knockout Consortium.

## Supplemental Material

**Figure S1:**
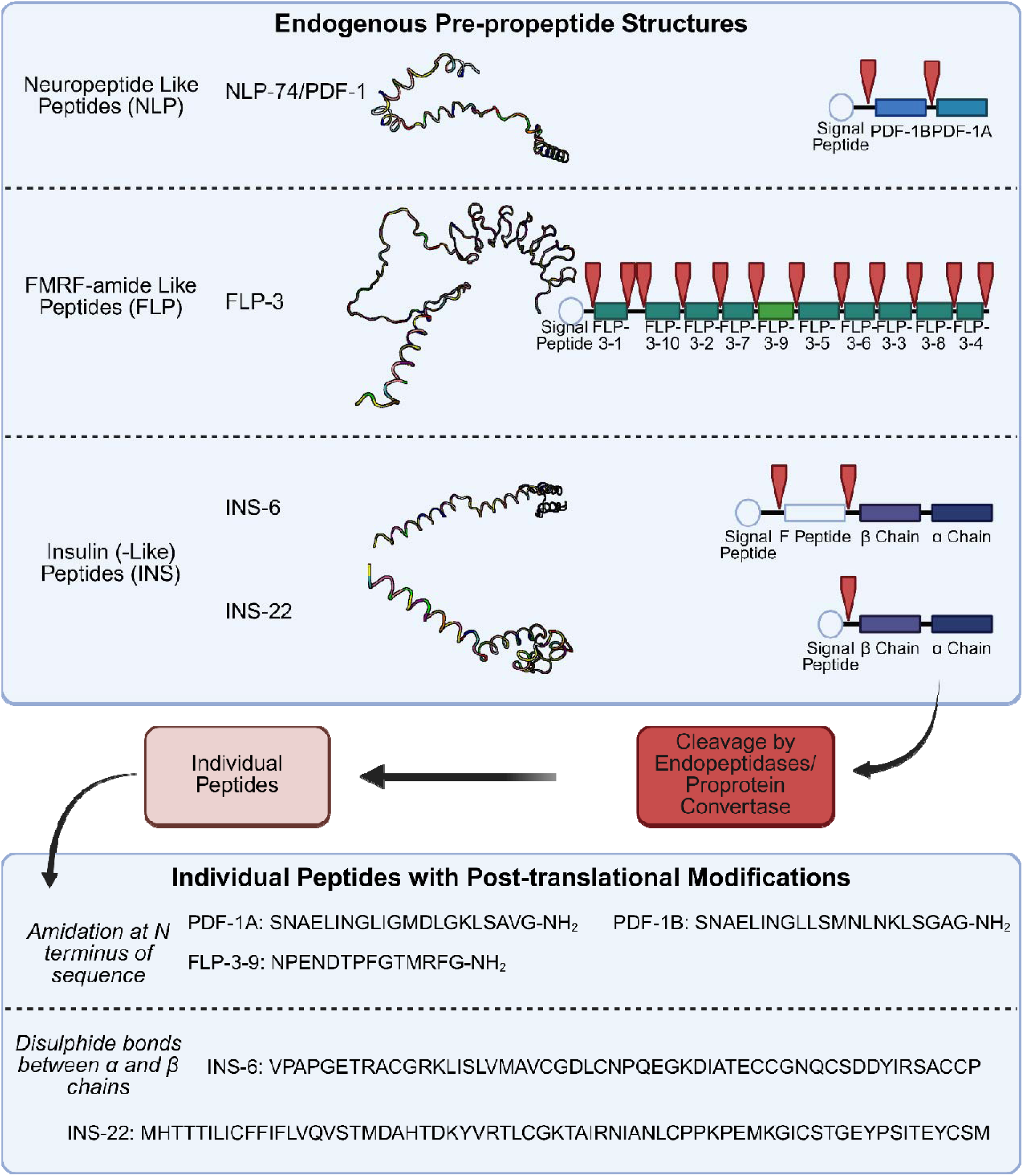
Endogenous Processing of Neuropeptides in *C. elegans*. Endogenous neuropeptide processing occurs inside neurons. From a neuropeptide precursor gene, an entire pre-propeptide is translated. We tested a representative peptide from each neuropeptide class in *C. elegans.* The DNA sequences for the neuropeptide precursor genes were from Wormbase.org and the putative unprocessed protein sequence and AlphaFold protein structure were obtained from uniprot.org. Once the pre-propeptide is translated, it is then processed by a series of enzymes. First the signal peptide is removed. And then the individual peptides are cleaved, at sites indicated by red arrows. We designed our neuropeptide constructs to contain these convertase enzyme cleavage sites, so we would anticipate that our delivered neuropeptide could be processed as demonstrated. Cleavage of individual peptides happens by the proprotein convertase enzymes. *C. elegans* have four protein convertases serving different functions in various tissues. Most common for neuropeptide processing appears to be EGL-3/KPC-2 (PC2 homolog cleaves K/R) but there are also, KPC-1 (cleaving K/R in dendrites), BLI-4/KPC-4 (cleaving K/R in the cuticle), and AEX-5/KPC-3 (cleaving paired basic amino acids in the muscle and intestine) (last three are PC1 homologs) [56, 101–103]. After protein cleavage creates individual neuropeptides, the peptides can be post-translationally modified. The FLP [68] and NLP peptides are often amidated [19] while INS peptides generally have disulfide bonds running between their α and β chains [58, 63]. Once the neuropeptides are processed, they are packaged into dense core vesicles for release at the synapse. They can be released with or without neurotransmitters. After binding to GPCRs on post-synaptic neurons, they slowly diffuse away from synapse and are broken down; this occurs at a slower rate than neurotransmitter reuptake leading to longer lasting neuropeptide effects [18, 104]. Created in BioRender. DiLoreto, E. (2025) https://BioRender.com/dyorxwg

**Figure S2:**
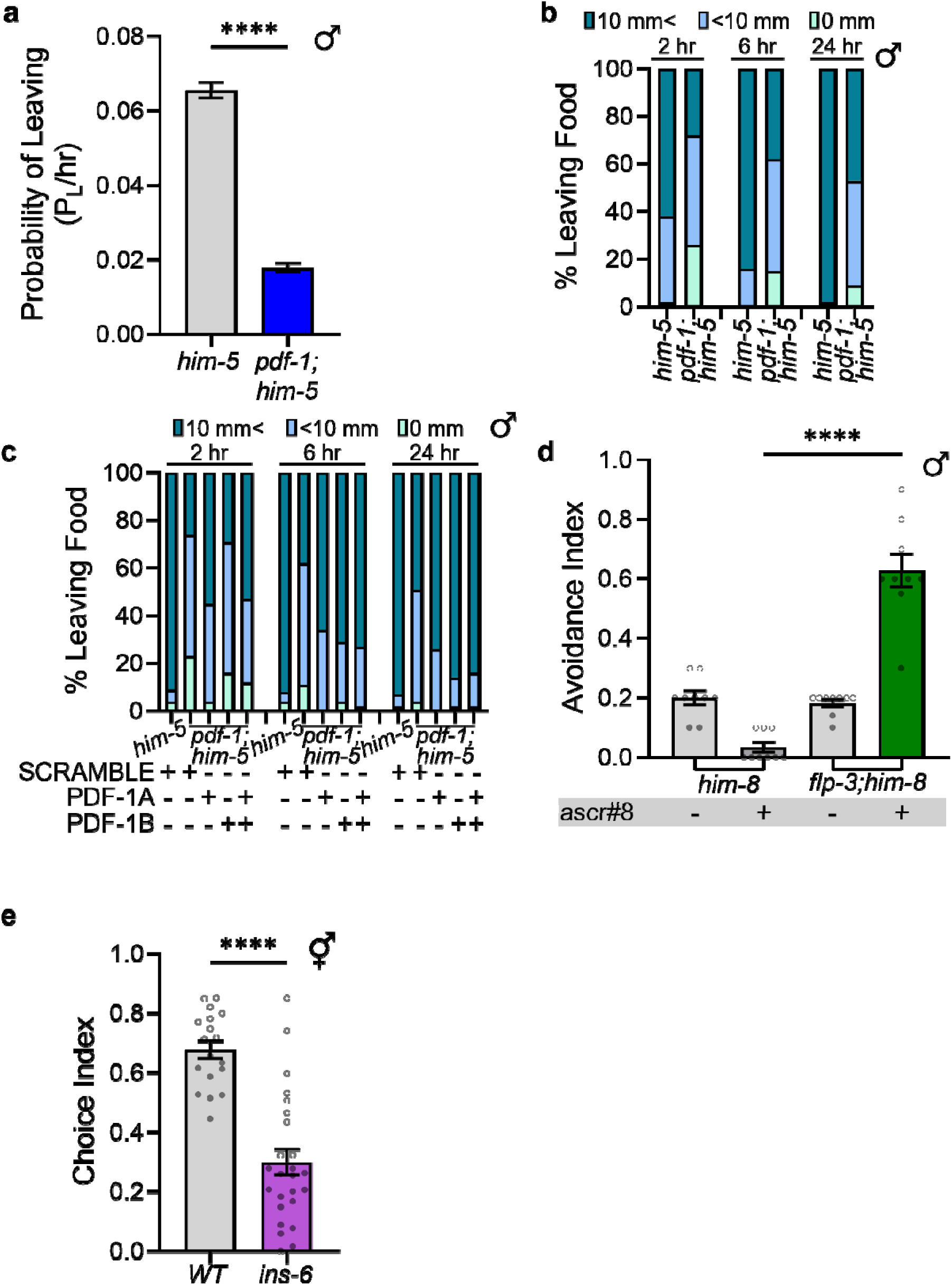
Neuropeptide gene deficient animals exhibit different behaviors than they wildtype counterparts. **A)** *pdf-1;him-5* males fed OP50 without peptide have a significantly lower rate of leaving food beyond 35 mm compared to wildtype *him-5* animals (****p ≤ 0.0001, unpaired t-test). **B)** The location of males can be traced across time and binned as animals who never left food (0 mm), those that had a minor excursion (10 mm >), and those with a major excursion (10 mm <). This is then converted to a percentage and plotted across time. These values contribute to the probability of leaving rate. *pdf-1* animals fed OP50 without peptide have lower rates of major excursions compared to wildtype *him-5* animals. **C)** SCRAMBLE DH5α *E. coli* fed *pdf-1 lof* males have a lower percentage of animals making major excursions off of food than *him-5* animals. **D)** Compared to wildtype *him-8* males, *flp-3;him-8 lof* males fed OP50 *E. coli* without peptide avoid ascr#8 at a significantly higher rate (****p ≤ 0.0001, Mann-Whitney test). **E)** Compared to wildtype animals, *ins-6* animals fed OP50 *E. coli* without peptide have a lower choice index towards high 750 mm NaCl (****p ≤ 0.0001, unpaired t-test).

**Figure S3:**
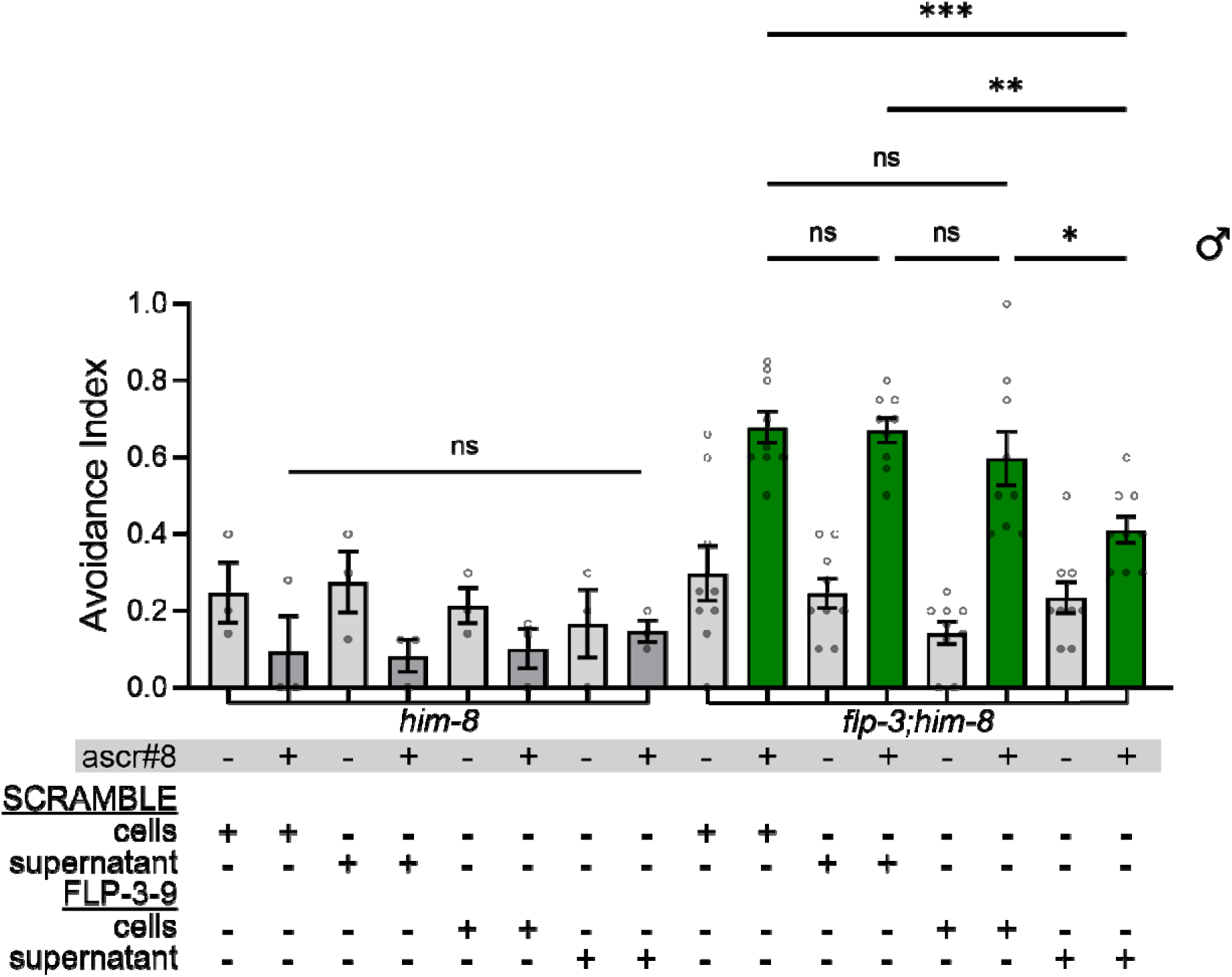
Bacterial cell supernatant from DH5α cells liquid culture growth alone may be sufficient to restore neuropeptide behavior. *him-8 lof* males fed the supernatant or cells derived from fed DH5α bacterial culture expressing the SCRAMBLE or FLP-3-9 peptide did not alter the lack of avoidance towards ascr#8. *flp-3;him-8* males fed the supernatant from a DH5α culture expressing FLP-3-9 did exhibit a suppression of avoidance behavior. (comparisons between ascr#8 stimulus (+) conditions within genotypes, ^ns^p ≥ 0.05, *p ≤ 0.05, **p ≤ 0.01,****p ≤ 0.0001, One-way ANOVA with Tukey’s multiple comparison)

**Figure S4:**
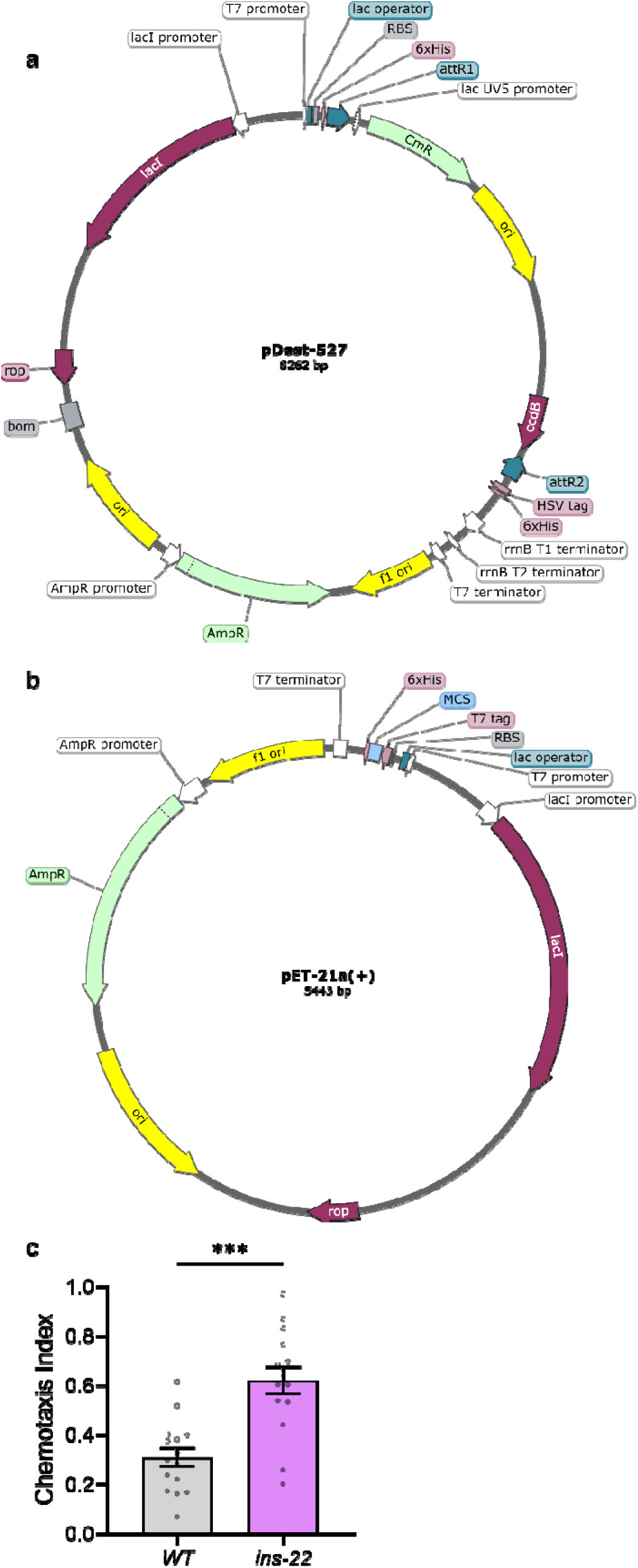
Plasmid maps of old Gateway cloning and new constructions. **A)** pDest-527 vector used in Gateway cloning. Created in SnapGene **B)** pET-21a(+) vector used in Genscript XhoI/XhoI construction. Created in SnapGene **C)** Chemotaxis values of wildtype (N2 Bristol) and *ins-22 lof* animals towards 10% butanone. (***p ≤ 0.001, Mann-Whitney test).

**Figure S5:**
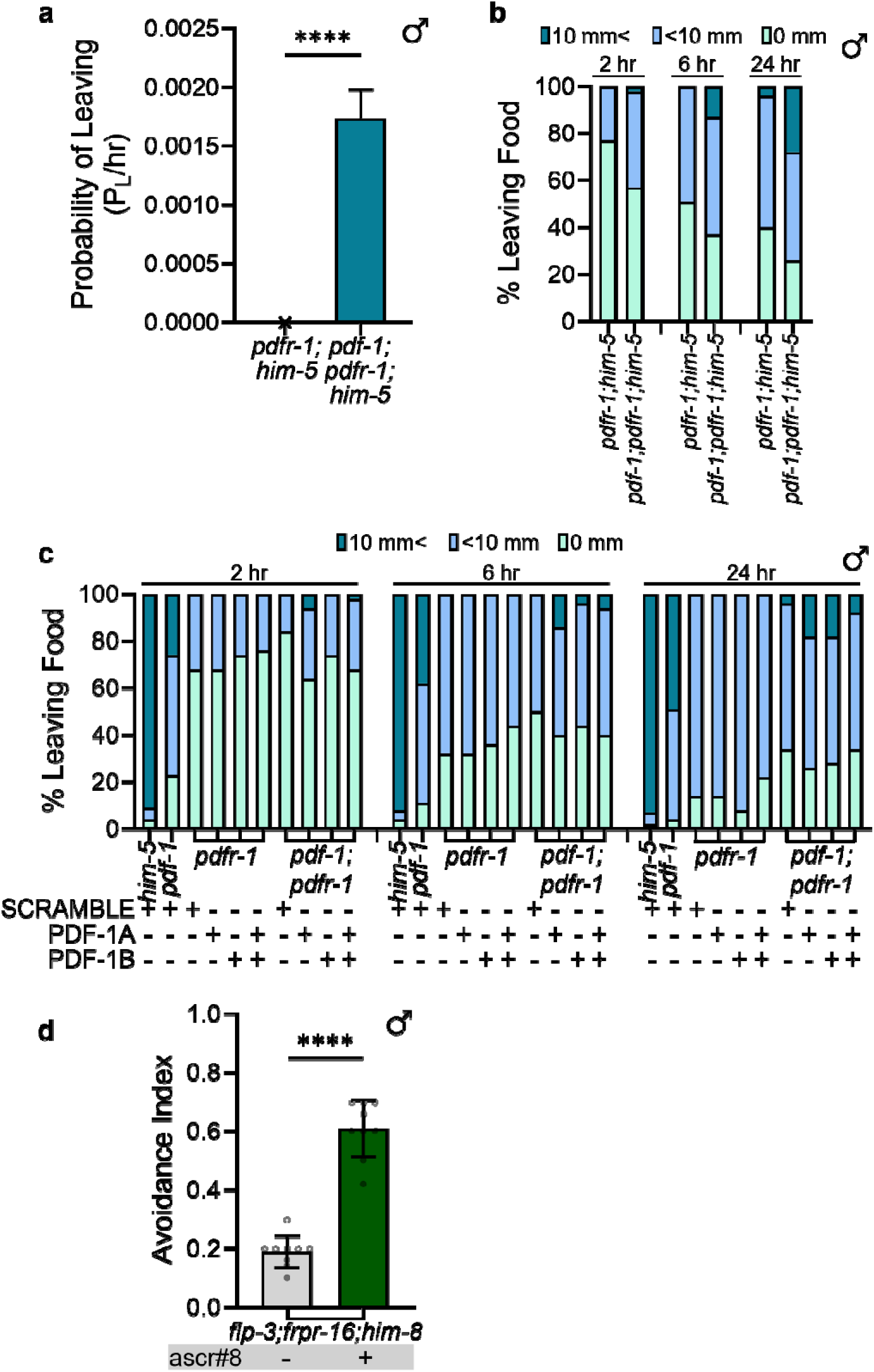
Neuropeptide receptor *lof* animals exhibit defective behavior without peptide feeding. **A)** *pdf-1;pdfr-1 lof* in a *him-5* background do leave food greater than 35 mm at a higher rate than *pdfr-1;him-5 lof* males (****p ≤ 0.0001, unpaired t-test). **B)** Excursion rates of neuropeptide receptor mutants (*pdfr-1;him-5*) and neuropeptide/neuropeptide receptor mutants (*pdf-1;pdfr-1;him-5*) fed OP50 *E. coli* do not readily leave food or have a major excursion 10 mm <. **C)** *pdf-1;him-5* neuropeptide receptor mutants fed PDF-1 peptides in DH5αa *E. coli* do not have excursions off food at high rates like *him-5* males. **D)** *flp-3;frpr-16* double mutants in a *him-8* background avoid ascr#8 at a higher rate than the solvent control (****p ≤ 0.0001, Mann-Whitney test).

**Figure S6:**
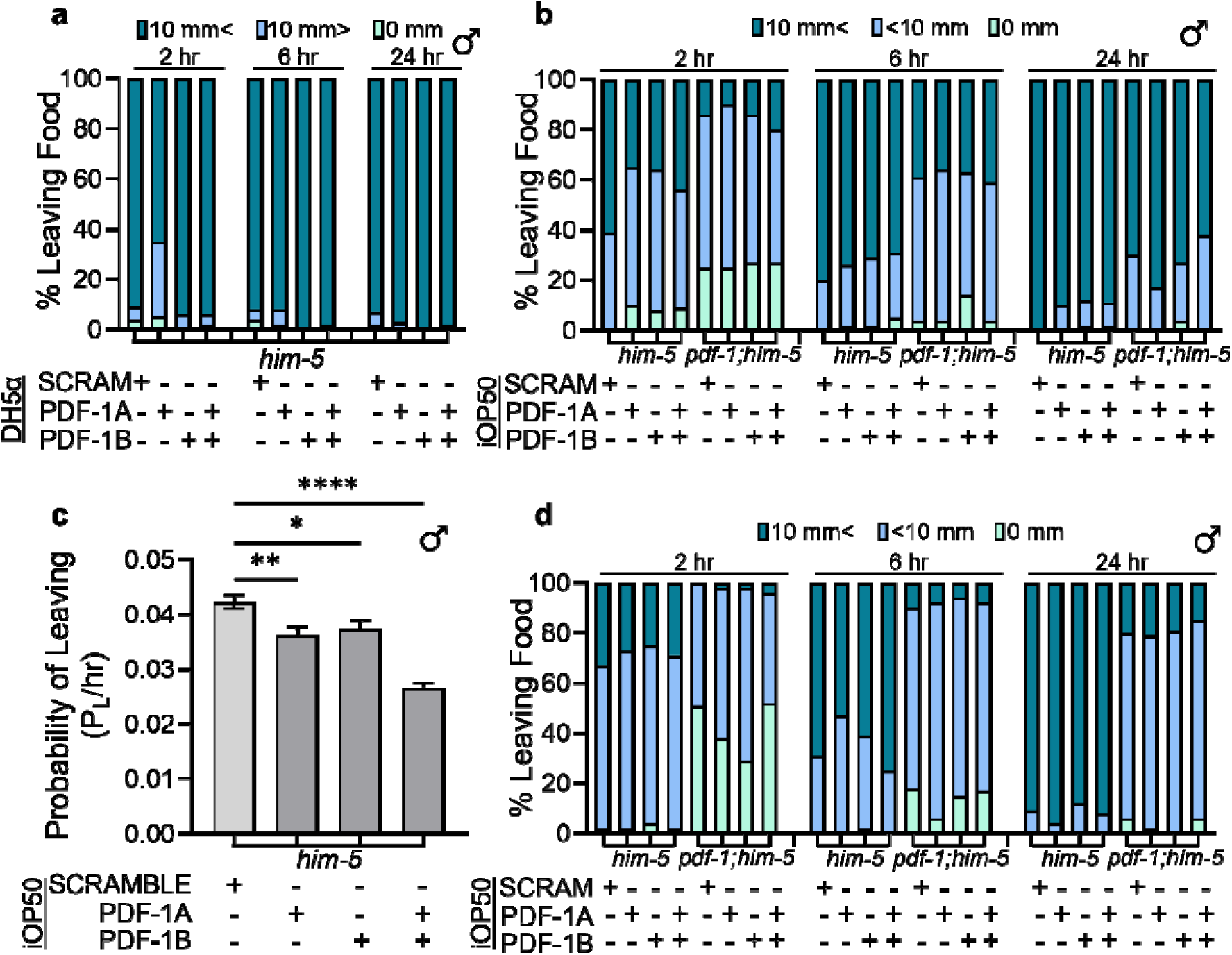
Overexpression of wildtype animals fed neuropeptide in DH5α or iOP50 *E. coli*. **A)** Most *him-5* males fed SCRAMBLE or PDF-1 peptides in DH5α *E. coli* made a major excursions traveling < 10 mm from food by 2 hours after plating. **B)** *him-5* males do not remain on food at the rates of *pdf-1;him-5* animals fed PDF-1 peptides in iOP50 *E. coli* and tested on iOP50 assay plates containing peptides. *pdf-1;him-5* males fed PDF-1 peptides in iOP50 *E. coli* have excursions off food at higher rates than SCRAMBLE-fed animals. **C)** *him-5* males fed PDF-1 peptides in iOP50 *E. coli* and then tested on OP50 assay plates not containing peptide, do not have excursions leaving food more than 35 mm at higher rates than SCRAMBLE fed males (*p ≤ 0.05, **p ≤ 0.01,****p ≤ 0.0001, One-way ANOVA with Dunnett’s post-hoc test). **D)** *him-5* males do not remain on food at the rates of *pdf-1;him-5* animals fed PDF-1 peptides in iOP50 *E. coli* and tested in OP50 assay plates not containing peptides.

**Figure S7:**
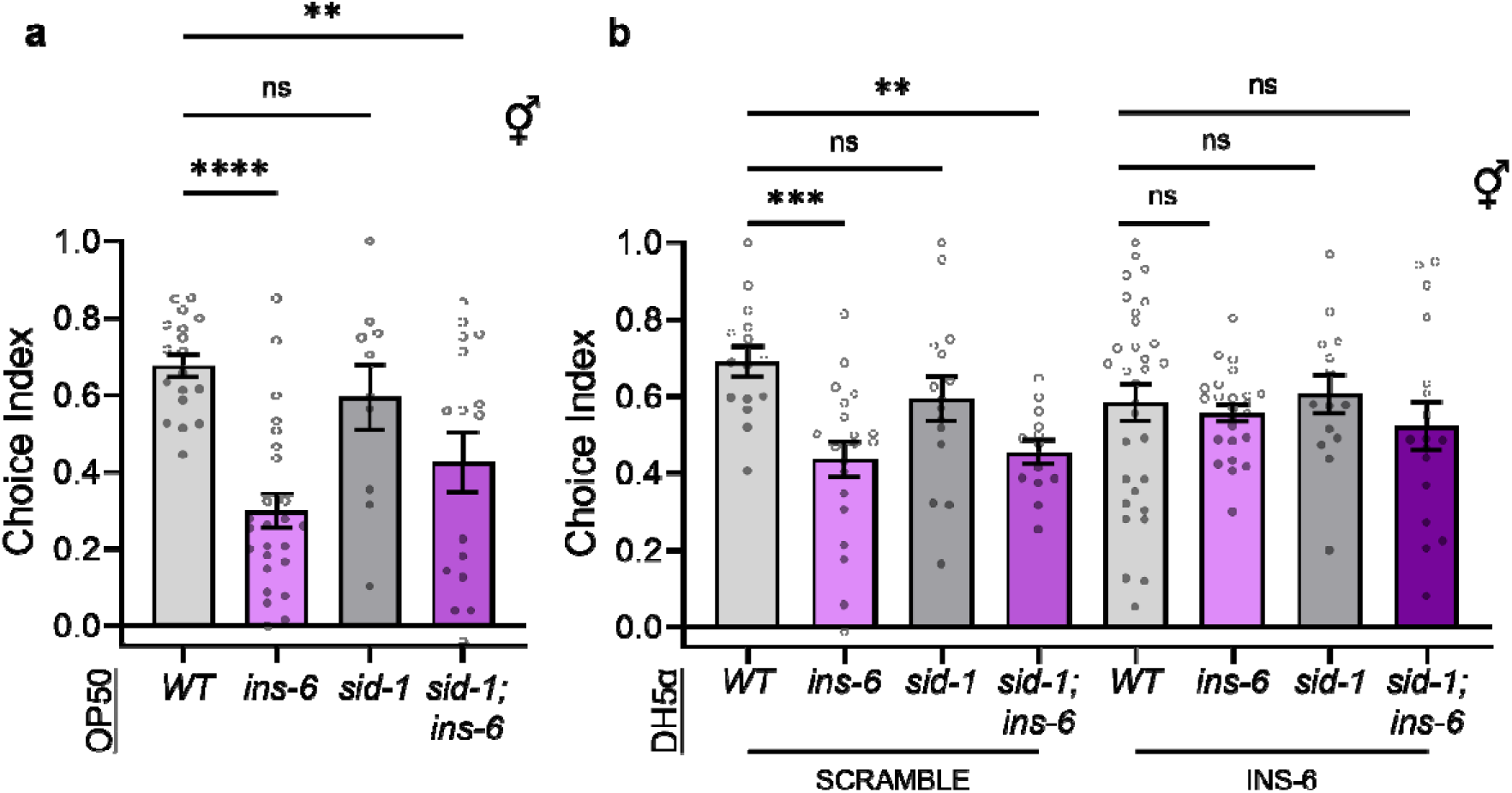
Investigating putative neuropeptide uptake mechanisms. **A)** Choice index of OP50 fed animals moving towards 0 or 750 mM NaCl. Mutants lacking *ins-6* showed reduced preference for 750 mM NaCl. (^ns^p ≥ 0.05, *p ≤ 0.05, **p ≤ 0.01,****p ≤ 0.0001, One-way ANOVA with Dunnett’s multiple comparison). **B)** Choice index of dsRNA transport mutant animals (*sid*-*1*) demonstrated that when fed DH5α bacteria expressing SCRAMBLE peptide both *ins-6 lof* and *sid-1;ins-6 lof* animals displayed defective NaCl attraction. After feeding of DH5α *E. coli* expressing INS-6, there was no statistically significant difference in the behavior towards NaCl compared to wildtype animals. (comparisons made within peptide feeding condition, ^ns^p ≥ 0.05, *p ≤ 0.05, **p ≤ 0.01,****p ≤ 0.0001, One-way ANOVA with Dunnett’s multiple comparison).

**Table S1:**
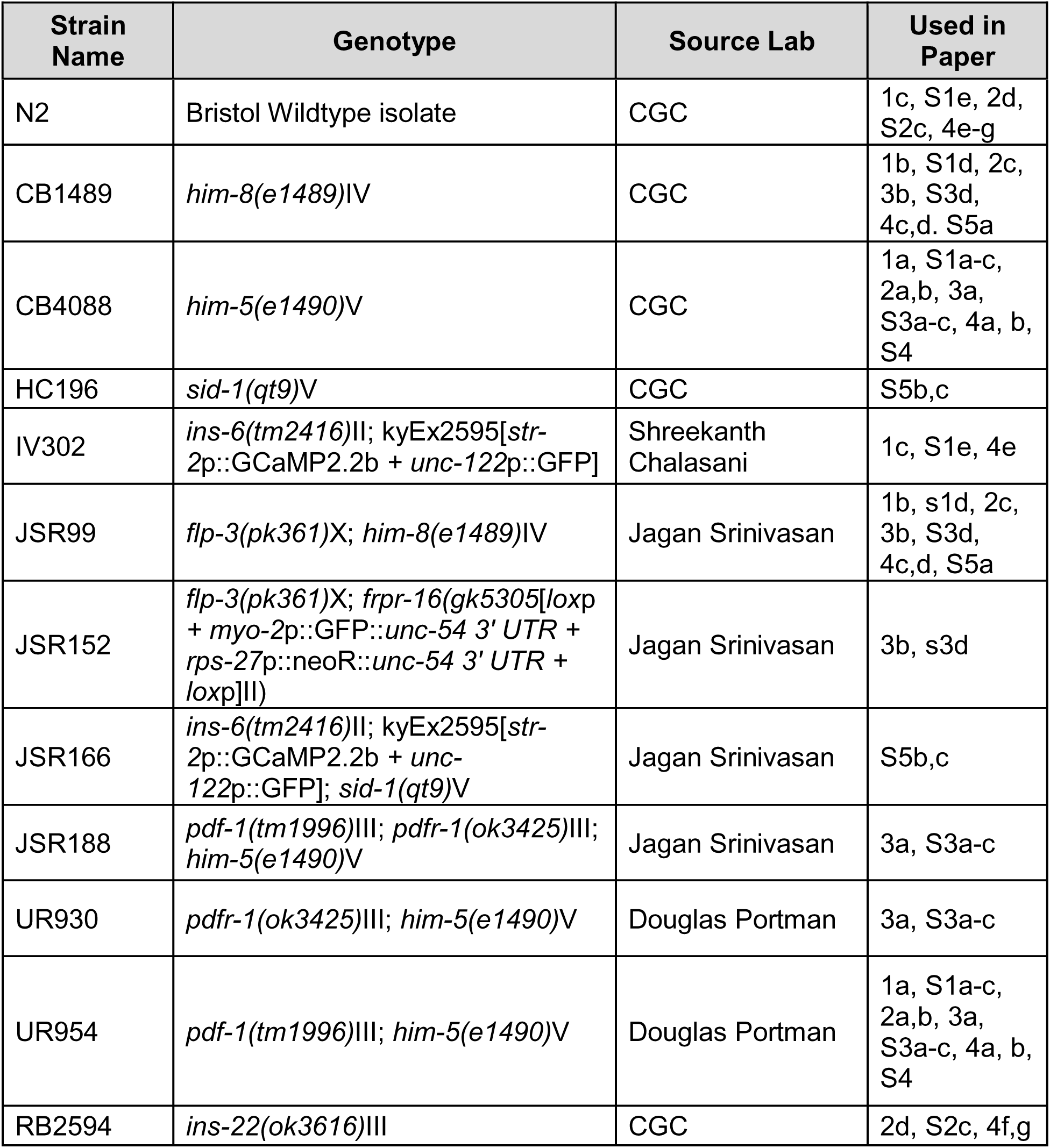
*C. elegans* Strains Used in Current Study.

**Table S2:**
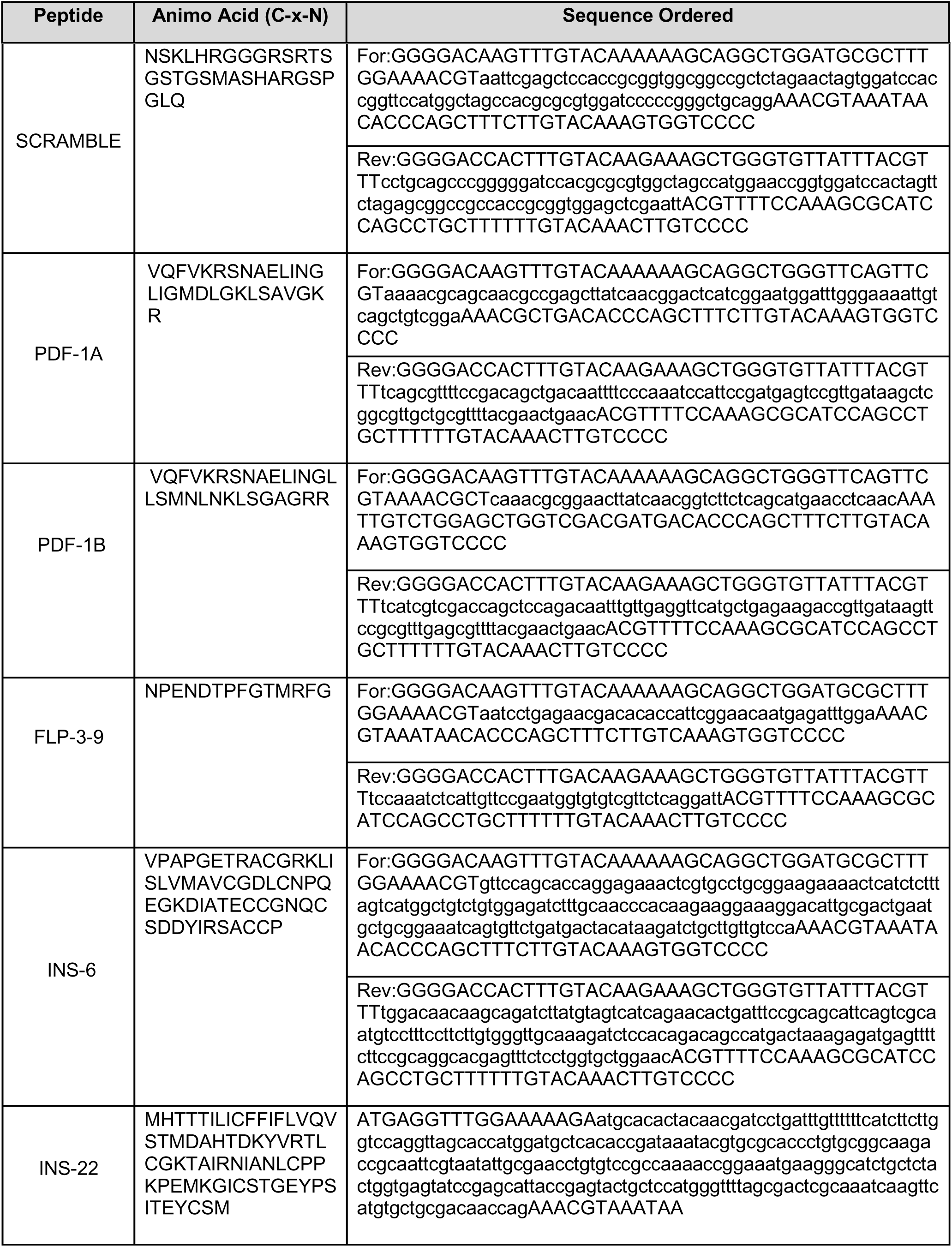
Neuropeptide Oligo Sequences Ordered for Individual Peptide Feeding. Neuropeptide DNA sequence indicated in lowercase.

